# Terahertz Circular Dichroism Spectroscopy of Biomaterials Enabled by Kirigami Polarization Modulators

**DOI:** 10.1101/549600

**Authors:** Won Jin Choi, Gong Cheng, Zhengyu Huang, Shuai Zhang, Theodore B. Norris, Nicholas A. Kotov

## Abstract

Terahertz circular dichroism (TCD) offers spectroscopic capabilities for understanding mesoscale chiral architecture and low-energy vibrations of macromolecules in (bio)materials^1–5^. However, the lack of dynamic polarization modulators comparable to polarization optics for other parts of electromagnetic spectrum impedes proliferation of TCD spectroscopy^6–10^. Here we show that tunable optical elements fabricated from patterned plasmonic sheets with periodic kirigami cuts make possible polarization modulation of THz radiation under application of mechanical strain. A herringbone pattern of microscale metal stripes enables dynamic range of polarization rotation modulation exceeding 80° over thousands of cycles. Upon out-of-plane buckling, the plasmonic stripes function as reconfigurable semi-helices of variable pitch aligned along the THz propagation direction. Several biomaterials, exemplified by elytrons of *Chrysina gloriosa* beetles, revealed distinct TCD fingerprints associated with the helical substructure in the biocomposite. Analogous kirigami modulators will also enable other applications in THz optics, such as polarization-based terahertz imaging and phase-encrypted telecommunication.

---

Scientific and technological advances related to chirality of liquid crystals, biomolecules, and synthetic drugs were enabled by the prior development of chiroptical spectroscopies, notably electronic circular dichroism (ECD) and vibrational circular dichroism (VCD), which enable identification of mirror asymmetry at molecular and nanometer scales. ECD and VCD are based on the modulation of circularly polarized light with photon energies in the ranges of 1.5-7 eV and 0.07-0.5 eV, respectively, which limit the physical dimensions and the resonant energies of the chiral structures that can be probed. Of particular interest is the far infrared (IR) or terahertz (THz) region of the electromagnetic spectrum corresponding to wavelengths in the range 0.1-1 mm and photon energies from ~0.001 eV to ~0.01 eV^11–15^ Besides being informative for many areas of THz studies from astronomy and solid-state physics to telecommunication, THz circular dichroism (TCD) is essential for understanding of biomaterials, biomolecules, and pharmaceuticals because the energy of THz photons enables probing the ‘soft’ oscillatory motions of biomolecules^1–5^. The practical realization of TCD, however, has proven to be an elusive goal due to the difficulties with polarization modulation of THz radiation. The key problem is the lack of optical components for modulation of circular polarization in the THz regime, which is easily accomplished at shorter wavelengths using photoelastic modulators (PEM), half- and quarter waveplates, and lately with chiral metamaterials and metasurfaces^6–9^ Although the modulation of linearly and circularly polarized THz beams has been demonstrated with fairly complicated and bulky optical systems based on THz metamaterials, e.g. with pneumatic control of scattering elements^6^, sufficiently strong and dynamic polarization rotation of THz radiation remains a significant challenge^6–10^.

Kirigami, the oriental art of paper cutting, presents a powerful tool to create complex and tunable three-dimensional (3D) geometries from simple (2D) two-dimensional cut patterns, which can be scaled across many orders of magnitude to yield macro to nanoscale structures^16–22^. The ability to achieve out-of-plane deformations and designed 3D shapes, the robustness of the patterns under cyclic reconfiguration and the manufacturing simplicity of kirigami structures together promise untapped possibilities for the efficient modulation of THz optical beams. Here we show that kirigami optics affords real-time modulation of THz beams with polarization rotation and ellipticity angles as large as 80° and 40° over thousands of cycles, respectively. The unusually large amplitudes of polarization rotation and ellipticity angles were enabled by double-scale patterns comprised of microscale metallic stripes together with wavelength-scale kirigami cuts.

The kirigami modulators in this study are made from parylene - a stiff polymer (Young’s modulus *E* = 2.8 GPa) with high transparency across the THz spectrum^23^. Parylene sheets were patterned with straight cuts in a face centered rectangular lattice with a periodicity of *p_cut_* = 600 μm (Supplementary Fig. 1 and 2). This 2D pattern transforms upon stretching into an array of alternating convex and concave out-of-plane surfaces due to buckling. (Fig. 1a and Supplementary Fig. 4).

**Figure 1.**
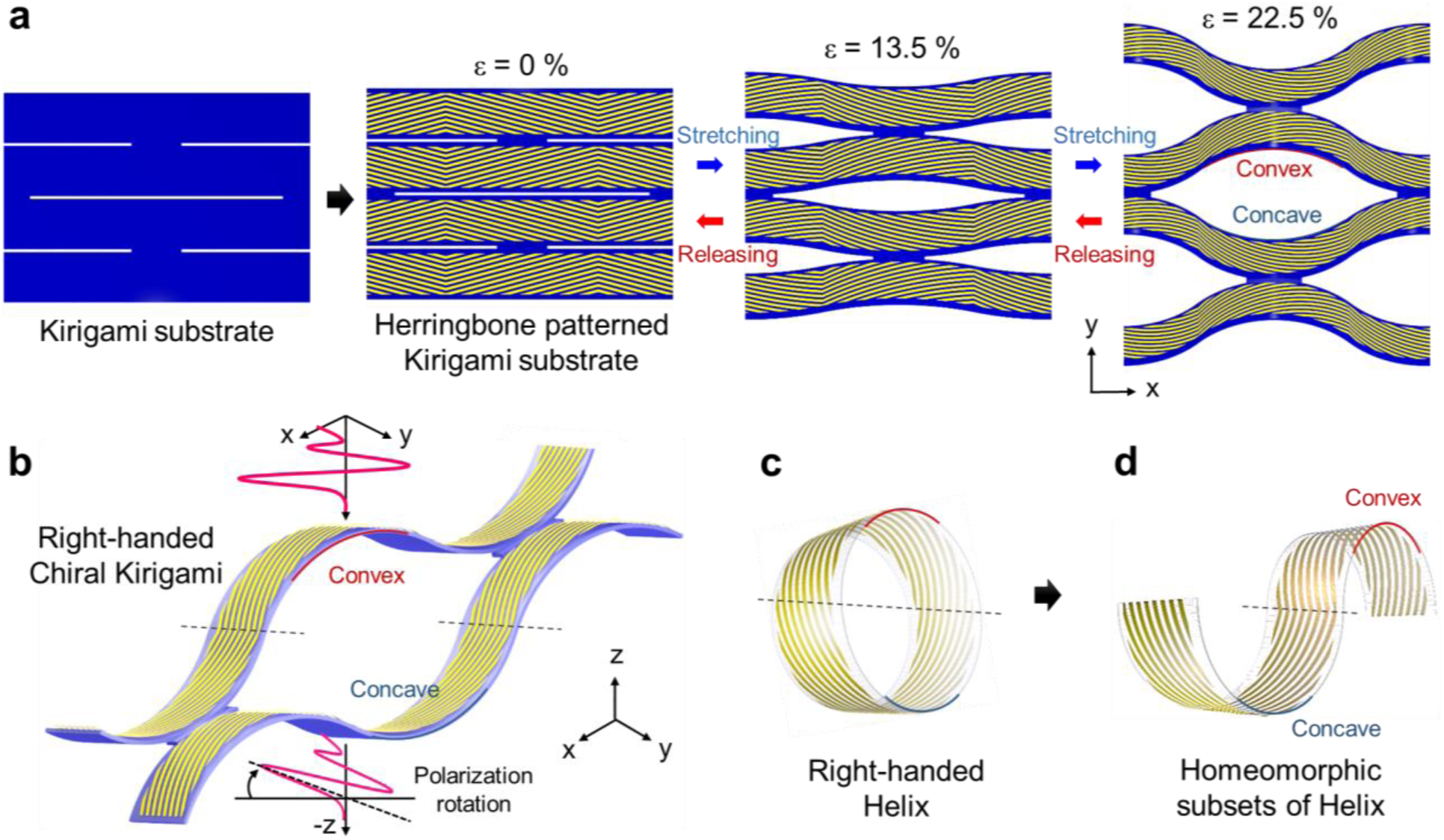
Schematic of chiral kirigami topology. **a,** Herringbone structured Au strips are deposited on the kirigami substrate. This chiral kirigami topology can tune the polarization rotation angle and ellipticity by mechanical force. **b,** Stretched chiral kirigami metamaterial that is topologically equivalent helix structure. **c,** Standard right-handed helix structure whose outside is covered with slanted striations and the structure which has homeomorphic subsets of helix.

The function-defining structural feature was gold herringbone pattern with *D_n_* symmetry deposited on parylene sheets in registry with kirigami cuts. When buckled, the patterned surface is transformed into a homeomorph of a three-dimensional helix (Fig. 1). Its pitch varies under mechanical strain while its long axis remains aligned with the surface normal (z-axis) and with propagating THz beam. Unlike microfabricated metallic helices^19,24^ and semi-dimensional metaoptics^25,26^, the double-patterns kirigami surfaces enable strong and tunable polarization rotation with real time modulation. Their confocal microscopy images (Supplementary Figs 4 and 5) obtained for strains *ε* from 0 % to 22.5 % demonstrated that buckling and tilting of each out-of-plane segment occurred simultaneously for the entire sheet, which is essential for uniform polarization front of a beam. The reconstructed contour maps of left-handed (*L-*) and right-handed (*R-*) kirigami structures (Fig. 2b, Supplementary Fig. 4) at *ε* = 22.5% strain indicate that the edges of the buckled elements extended to 68 μm symmetrically along the positive and negative z-axis, tuning the radius and pitch of the clockwise and counter clockwise half-helices ^19,24^ are the two key factors for controlling electrodynamic interactions of these structures with left- and right circularly polarized photons. The experimental deformations matched the predictions from finite-element modeling exactly (Supplementary Fig. 4c and 4d).

**Figure 2.**
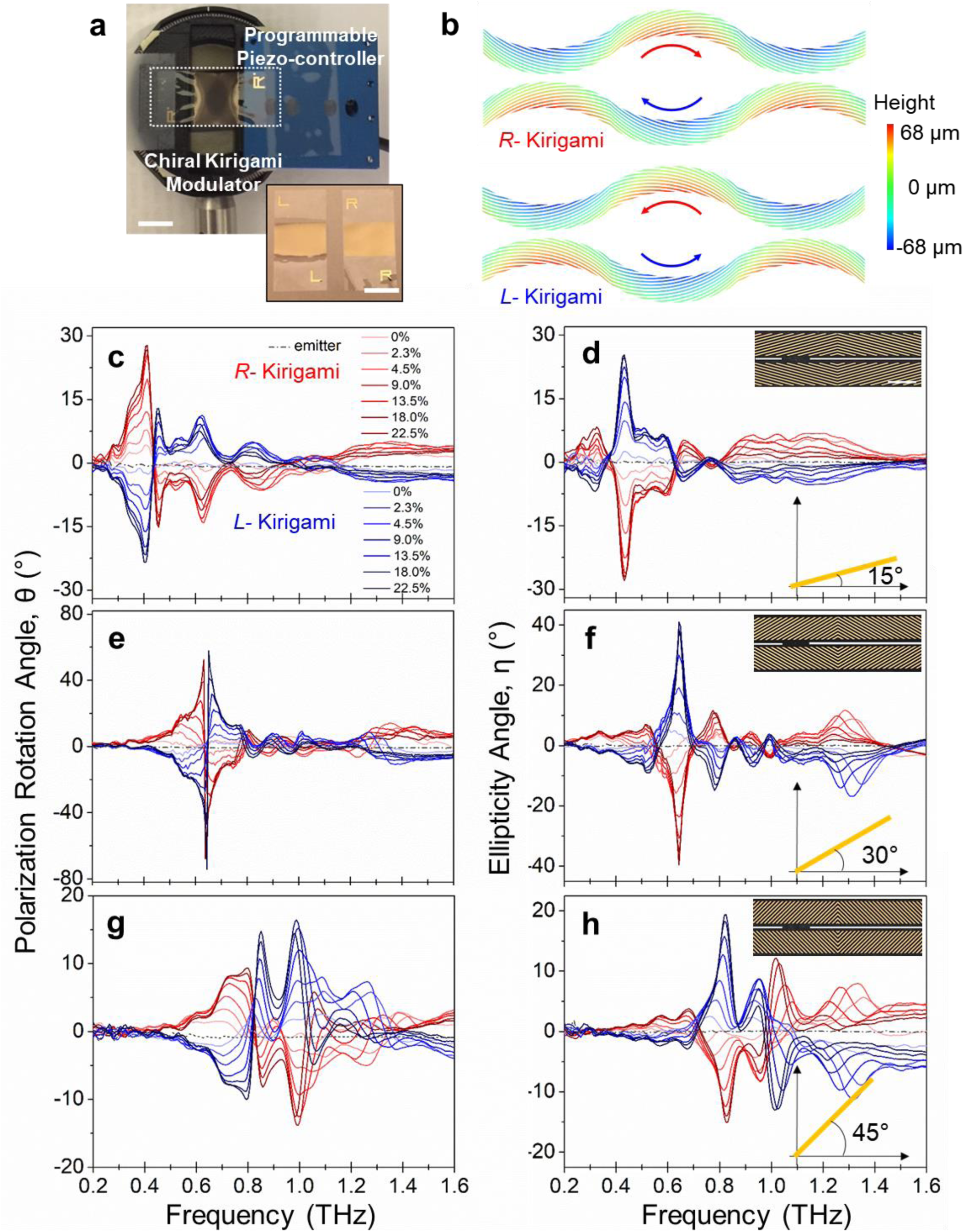
THz-TDS measurement of chiral kirigami modulator. **a,** kirigami mounted on the 3D printed rotatable optical holder with piezoelectric controller. Subset figure shows photo image of left and right handed chiral kirigami substrate. Yellow shiny region is the herringbone patterned Au zone. Both of scale bars are 1 cm. **b,** Contour map of kirigami modulator reconstructed from laser confocal microscope images. **c, e and g,** Results of polarization rotation angle of kirigami having slanted Au angles (*φ*) of 15, 30, 45 degree with respect to strain (%), respectively. **d, f and h,** Results of consequent ellipticity angle of kirigami having slanted Au angles (*φ*) of 15, 30, 45 degree with respect to various strain, respectively. One can notice that there is an approximately 0.2 THz increment per 15° change of *φ*. As can be expected by the Kramers-Kronig relation^11^, the ellipticity exhibited a dispersive curve and crossed zero at slightly off-resonance frequencies, where the polarization rotation showed maximum. Inset shows optical microscope images of each kirigami samples. Scale bar in **d** is 100 μm.

THz time-domain spectroscopy (THz-TDS) over the range 0.2-2 THz was used to characterize chiroptical performance of the kirigami modulators. The polarization rotation angle, *θ*, and ellipticity angle *ƞ* (Supplementary Information) of the THz beam after passing through kirigami sheets or biomaterials with expected TCD activity were determined using two complementary methods. The first protocol was based on standard calculations of Stokes parameters from the Jones matrix from a sequence of linear polarization measurements; the second one was based on direct measurements employing kirigami modulators.

The kirigami sheets were mounted on an optical holder and finely controllable stress was applied with a programmable piezoactuator with a precision of 100 nm (*ε* = 0.001%) (Fig. 2a). As expected *θ* and *ƞ* increased with strain and kirigami structures with left-handed and right-handed herringbone patterns exhibit THz responses that are nearly identical but with opposite signs (Fig. 2c-2h). The inclination angle (*φ*) of the herringbone patterns (insets of Fig. 2d, f, h) determined the position of the main resonance peaks, which were observed at 0.41 THz for *φ* = 15°, 0.62 THz for *φ* = 30° and 0.81 THz for *φ* = 45°. The maximum values of *θ* and *ƞ* reached as high as 80° and 40°, respectively, were obtained for herringbone patterns with *φ* of 30°. This maximum ellipticity value is almost close to that of quarter-waveplate. Note that the magnitudes of *θ* and *ƞ* can be different depending on the in-plane rotation angles due to birefringence, which was taken into account in the TCD spectra (Supplementary Figs 8-11). As a control, an achiral pattern with horizontally aligned (*φ* = 0°) Au strips was tested, and showed near-zero values of *θ* and *ƞ* regardless of the strain, confirming the critical role of the double-pattern design for the strong optical activity (Supplementary Fig. 11). Polarization modulations with nearly identical values of *θ* and *ƞ* were obtained for 1000 cycles with *ε* between 0 % and 22.5 % (Supplementary Fig. 12).

The effect of the microscale cut pattern on the optical performance of the kirigami polarization modulators was tested for variable size of the unit cell for a constant inclination *φ* = 30°. As *L_cut_* becomes larger, the main TCD peak shifts to the red (Fig. 3a and 3b), indicating that its spectral position is determined by the longitudinal plasmonic resonances of the Au strips. Confirming this conclusion, TCD spectra of kirigami modulators with Au strips having same length and *φ* but narrower width display the same position of resonance peak and similar overall shape (Supplementary Fig. 11). The resonance wavelength of the kirigami sheets with herringbone patterns can be heuristically assessed as an *LC* circuit with the resonance frequency of 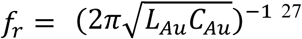. The inductance *L_Au_* and capacitance *C_Au_* of Au strips scale linearly with its length, *l* (Fig. 3c, Supporting Information), and therefore *f_r_* becomes inversely proportional to *l* (Fig. 3d). Alternatively, the metal strips can also be approximated as Hertzian dipoles bent and tilted in 3D space, *l* ~ λ_*r*_ = *c*/4*f*_*r*_, where *c* and λ_*r*_ are the speed of light and resonance wavelength. This equation can be used to provide an approximate guide of the design of herringbone patterns for different applications.

**Figure 3.**
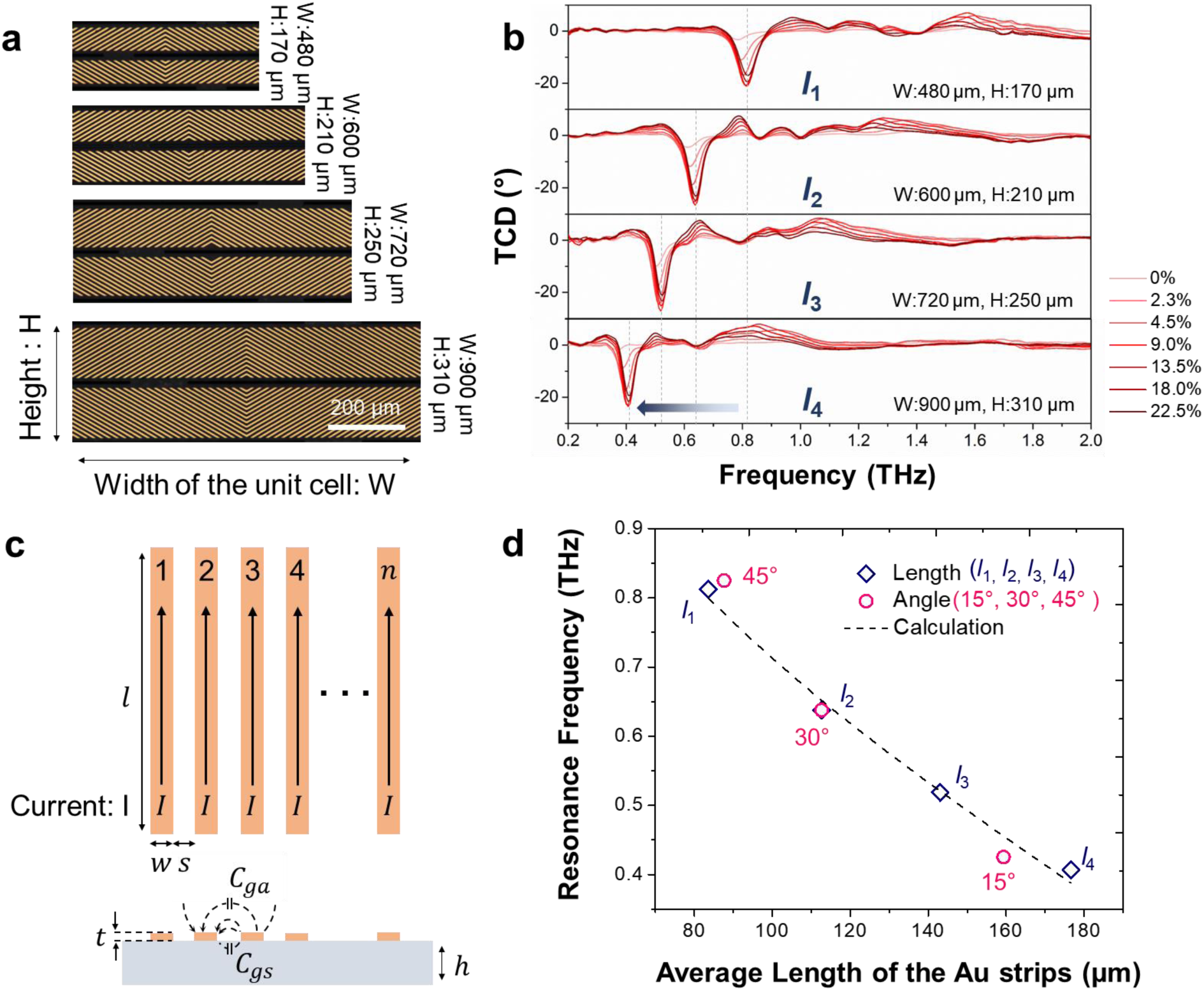
Understanding physical meaning of resonance frequency. **a,** Optical microscope image of various length of unit cell. All samples have *φ* of 30°. **b,** Result of measured TCD from *R-* kirigami modulator with various length of unit cell. **c,** Upper figure shows parallel conducting strips to obtain total inductance of this configuration. Lower figure is for calculating capacitance of array of strips. **d,** Relation between resonance frequency and average length (*l*) of the Au strips. Scale bar in **a** is 200 μm.

TCD spectra of the kirigami optical components can be predicted with *ab ovo* electrodynamic simulations. Computed TCD spectra (Fig. 4a) matched well the experimental data with respect to the signs of the polarization rotation angle, peak positions, relative peak widths and amplitudes (Supplementary Fig. 11a). Calculated time-averaged current norm distributions generated on the Au strips for the incident of the circularly polarized beam point to the origin of the plasmonic states responsible for individual peaks (Fig. 4b-d)^6^. At the off-resonant frequency of 0.57 THz, the induced currents are low for both co- and cross-circularly polarized beam and most of the Au strips are optically inactive (Fig. 4b). At the resonant frequency of 0.82 THz, however, the incident beam induces strong currents in the Au strips. Simultaneously, the currents excited by the right-handed circularly polarized beam significantly exceed those for left circularly polarized beam (Fig. 4c and 4d). As a result, the transmittance of the left circularly polarized beam is larger than that of the right circularly polarized beam due to the induced current, which is consistent with the sign of the peaks in Fig. 4a and Supplementary Fig. 11a.

**Figure 4.**
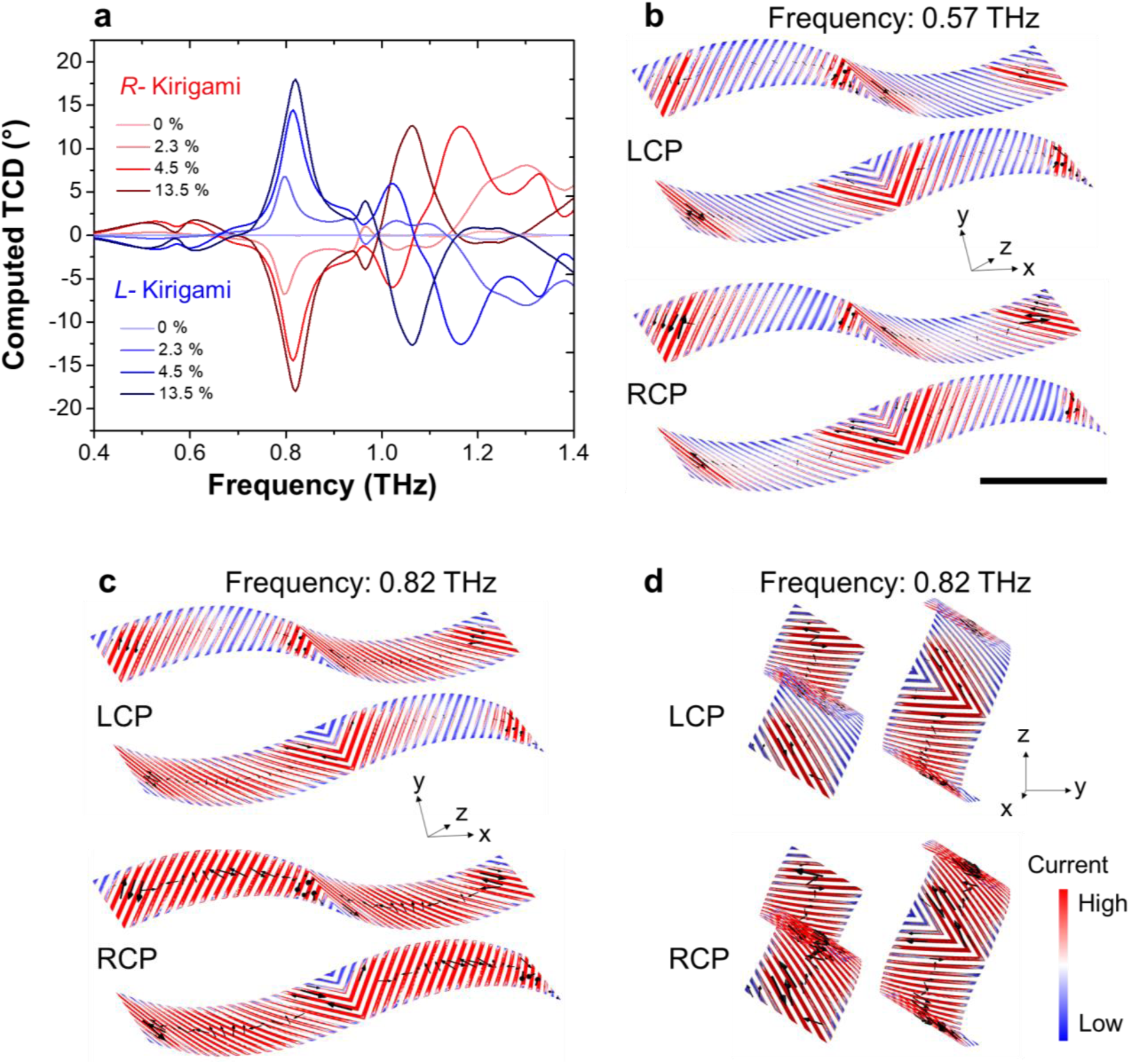
Computed terahertz circular dichroism and time-averaged current norm distributions on kirigami modulator (*φ*=45°). **a,** Computed TCD spectrum for 4 deformation states. **b and c,** Tilted view of current norm distributions of *R-* kirigami at the frequency of 0.57 THz, 0.82THz, respectively. **d,** Side view of current norm distributions of *R-* kirigami at 0.82 THz. Black arrows indicate the current directions. A scale bar is 200 μm.

The unique combination of high ellipticity and tunability of kirigami optics makes possible its utilization of for modulating THz light beams in practical realizations of TCD spectroscopy to investigate biological and other materials that are opaque in the visible range but transparent for THz radiation. To demonstrate this capability, we measured TCD spectra of several representative biological samples (Fig. 5a and Supplementary Fig. 22), including a leaf of sugar maple tree (*Acer saccharum*), an elytron of green beetle (*Chrysina gloriosa*), a petal of dandelion (*Taraxacum officinale*) and a piece of pig fat. Here, TCD spectra were calculated directly from difference of transmission-intensity between left and right elliptically polarized THz beam (EPB) generated by kirigami modulators according to Eq. 1:

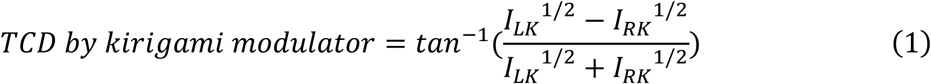

where, *I_LK_* and *I_RK_* are the intensities of the left and right EPB after passing through sample, respectively. We observed distinct THz spectra from the tested biomaterials that can be associated with low energy vibrational modes of their biological components and with chiral structural organization. In both cases, the opaqueness of biomaterial and mismatch in energy/wavelengths with visible light would not allow chiral characterization by ECD or VCD spectroscopy. An exemplary case is the transmissive TCD measurements of an elytron of *C. gloriosa* beetle (Fig.5b and c), which is known to have the selective reflection of circularly polarized light in the visible range^28^. A positive peak of TCD (Fig.5g) as large as about 3° at 0.68 THz is observed in the red circled area in Fig.5f. Notably, the absorption peak of this biological composite (Fig.5h) is well aligned with that of TCD peak. This TCD spectrum is associated with the chirality of its micro structure of exocuticle (Fig. 5d) which shows chiroptical response in reflected light in visible range (Fig. 5e)^28^.

**Figure 5.**
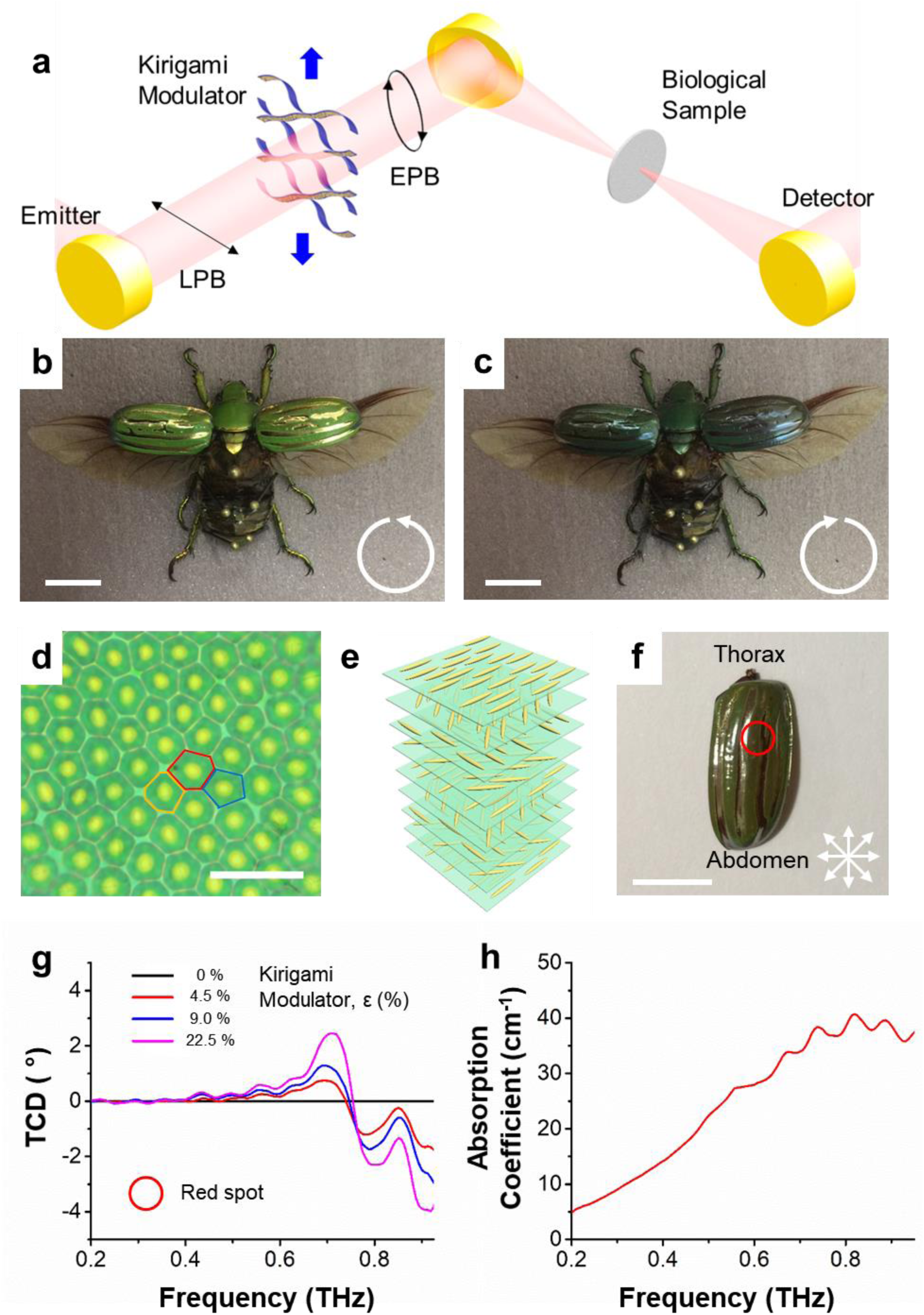
Measurements of TCD using kirigami chiroptical modulator. **a,** Schematic of TCD spectroscopy using kirigami modulator. A focused THz beam with ~500 μm spot size was used to explore biological sample. LPB and EPB indicate the linearly and elliptically polarized beam in respectively. **b and c,** Photographs of the beetle *C.gloriosa* with a left and right circular polarizer front of the camera, respectively. **d,** An optical microscopy image of the exoskeleton of beetle *C.gloriosa*. The shape of the cells is pentagonal in blue, hexagonal in red and heptagonal in orange. Scale bar is 20 μm. **e,** Schematic representation of Bouligand structure. **f,** Image of an elytron of *C.gloriosa* without polarizer. Red circle indicates the spot corresponding to the TCD measurements. **g,** TCD spectrum from *C.gloriosa* measured by kirigami modulator at four different strains (%). **h,** Measured absorption coefficient of *C.gloriosa.* Scale bars in **b**, **c and f** are 1 cm.

In conclusion, the double-pattern design of kirigami materials combining submillimeter cuts and nanometer scale plasmonic stripes affords the real-time tunability of helical structures oriented perpendicularly to the propagation of the light beam. Kirigami optical elements make possible realization of TCD spectroscopy and better understanding of liquid-crystal-like organization of soft and mineralized tissues ^29,30^. The lightweight capabilities and high polarization efficiency of kirigami optics open a possibility of portable compact THz spectrometers. The realization of real-time polarization modulation of THz beams also enable advances in secure high bandwidth communication and non-invasive imaging.

