## Supplemental material for "Terahertz Circular Dichroism Spectroscopy of Biomaterials Enabled by Kirigami Polarization Modulators"

### Chiroptical Kirigami Modulators for Terahertz Circular Dichroism Spectroscopy of Biomaterials

**This PDF file includes:**

Materials and Methods
Figs. S1 to S23
Captions for Movies S1 to S2
References

**Other Supplementary Materials for this manuscript includes the following:**

Movies S1 to S2

### Table of Contents

**Materials and fabrications**
- Fabrication of kirigami modulators…………………………………………………….……3
- Dimension of kirigami pattern and definition of slant angle ($\varphi$)……………………………3
- Integration kirigami sheets with piezo-controller……………………………………………4

**Mechanical and cycling characterization**
- Mechanical and cycling characterization of kirigami sheets………………………………...4
- Finite-element modeling for mechanical characterization…………………………………...5

**THz-TDS measurements**
- THz-TDS measurements……………………………………………………………………..5
- Definition of sample mounting orientations and abbreviations for sample configurations….6
- Calculations of transmittance, polarization state and THz circular dichroism (TCD)……….6
- An example of raw THz-TDS data for kirigami sheets……………………………………...10
- Experimental data of transmittance, polarization state and TCD……………………………11

**Modeling of kirigami geometries and TCD simulations**
- Measurements and modeling of kirigami 3D geometries……………………………………11
- TCD simulation and surface current norm distributions……………………………………..11

**LC circuit model for frequency resonance**
- Circuit model for calculating resonance frequency…………………………………………..13

**TCD measured by kirigami modulator**
- Experimental setup and data of polarization states for stacked kirigami modulators………...14
- TCD of kirigami sample using kirigami modulator...............................................................15
- TCD of biological samples using kirigami modulator………………………………………..16

Fabrication of kirigami modulators

Poly(methyl methacrylate) (PMMA 950 C4, Microchem) is spin-coated (3000 rpm) on a 4 in. silicon wafer as a sacrificial layer and baked subsequently on a 180° hot plate to dry. Parylene C (SCS Inc.) is deposited on the PMMA-coated silicon wafer by a chemical vapour deposition system (PDS 2035CR, SCS Inc.). The thickness of Parylene C is set to ~6 μm and confirmed by surface profiler (Dektak XT, Bruker) after deposition. Herringbone patterned thin layers of chromium (~5 nm) and gold (~45 nm) are deposited on the Parylene C using electron beam evaporator (Enerjet evaporator) after a standard photolithography processes (MA/BA6 Mask/Bond aligner, Suss Microtec). Kirigami patterns are generated by additional photolithography on top of herringbone patterned substrate and followed by deposition of aluminum (~70 nm) as a masking layer for the reactive ion etching process. The corresponding patterns are formed by reactive ion etching (LAM 9400, Lam Research) of Parylene C. Lastly, the whole patterned wafer is soaked in aluminum etchant solution (Aluminum etch Type A, Transene) to remove the aluminum masking layer and in acetone to dissolve the PMMA sacrificial layer. The released kirigami sheet is rinsed carefully with isopropyl alcohol and distilled water and dried. The schematic is shown in Figure S1 to clarify each step.

Dimension of kirigami pattern and definition of slant angle ($\varphi$)

Fig. S2A shows a top view of the kirigami cut pattern without Au strips to exhibit the dimensions of the cut pattern. The length ($L_{cut}$) and height ($H_{cut}$) of each cut is 500 μm and 5 μm, respectively. The horizontal and vertical spacings between cuts are set to 100 μm resulting in a horizontal period of 600 μm and a vertical period of 105 μm. A detailed view of a single unit of slanted Au strips is shown in Fig. S2B. The width of each Au strip is set to 5 μm. The width and height of the total domain are 300 and 80 μm, respectively. Here, the slant angle ($\varphi$) is defined as the angle between the cut direction and the longer axial direction of the Au strips, as shown in Figure S2B. In this work, we tested kirigami samples with four different angles $\varphi$: 15°, 30°, 37.5° and 45°. Fig. S2C shows a top view of the double pattern kirigami, that is, the kirigami cut pattern together with the Au herringbone pattern. The red box indicates the unit cell of the extended modulator structure. The width ($W_{unit}$) and height ($H_{unit}$) of unit cell are 600 and 210 μm, respectively.

Integration kirigami sheets with piezo-controller

Fabrication of the chiral kirigami modulators is completed by integrating the kirigami sheet with a piezo-controller for the application of controlled strain ($\varepsilon$). We used a U-521 PI Line (PI instrument) linear position stage with 3D printed sample holder. For the stacked configuration used in terahertz circular dichroism (TCD) measurements, we used two U-521 piezo-controllers and manipulated them individually. This piezo-controller can be programmed with very high spatial precision of 0.1 μm ($\varepsilon$ = 0.001%). The applied strain values of 2.3, 4.5, 9.0, 13.5, 18, 22.5 % in the measurements presented in the main text and supporting materials is calculated from stretching distances of 0.2, 0.4, 0.8, 1.2, 1.6, 2.0 mm, respectively.

Mechanical and cycling characterization of kirigami sheets

The high elasticity and tunability of kirigami sheets are significant advantages of kirigami chiroptical modulators. This is because a network of notches made in a rigid substrate greatly increases the ultimate strain that can be applied to the sheets and prevents unpredictable local failure (*31*). Uniaxial tensile tests were performed by means of a TA.XT *plus* Texture Analyzer (Texture Technologies) and the *Exponent* (Texture Technologies) software package for tensile and cycling tests with a 0.5 N load cell at a constant strain rate of 0.2% per second. The engineering stress-strain data were obtained and each curve was averaged over 5 samples. Our kirigami modulators were found to reach strains as high as 150% without failure (Fig. S3A). Before cutting, the pristine parylene sheets shows a strain of 3.8%. In contrast, a kirigami cut significantly modifies the deformation behavior of a sheet, resulting in a lower stiffness and higher elongation than a pristine sheet, as seen in Fig. S3B. In the stress-strain curves, the kirigami sheet's initial state at < 4% strain is elastic in-plane deformation (Fig. S3B, blue section). As the applied stress exceeds a critical strain, the domains of the kirigami structure start to deform elastically in out-of-plane directions. Within this region (Fig. S3B, pink section), buckling occurs as the domains rotate to align with the direction of tensile stress as shown in Fig S4 and S5. After that, plastic deformation occurs and finally failure begins when one of the cuts begins to tear and crease (Fig. S3B, white section) (*31*). Since we only apply strains up to 22.5% to achieve out-of-plane deformations with convex and concave domains, the deformation is completely within the elastic region. Fig. S3C shows the stress-strain curve after 10000$^{th}$ cycle of stretching and releasing and surprisingly it is nearly identical to its initial curve.

Finite-element modeling for mechanical characterization

Commercial finite-element software (COMSOL Multiphysics 5.2a, COMSOL Inc.) was used to explore the strain distribution in kirigami sheets, which yields insight into the basic mechanisms governing deformation behavior. An approximate global mesh size of 25 μm was used. We set the following boundary condition on each side of the kirigami sheet in the axial direction: 1) at one end we fix it and no displacement is allowed to this boundary; 2) at the opposite end we enforce a load in the axial direction. Since, in real systems, there is always an asymmetrical force, we apply a very small bias force (approximately $10^{-4}$ times smaller than the load) on top edge of each cut and then pull in the axial direction (*31*). The FEM shows that high elasticity is due to the even distribution of stress over the kirigami sheet rather than concentrating on singularities (Fig. S4D and E) (*31*).

THz-TDS measurements

Terahertz time-domain spectroscopy (THz-TDS) was used to measure the optical responses of the chiral kirigami modulators. A Ti:Sapphire regenerative amplifier (RegA 9050, Coherent) with a center wavelength of 800 nm, a pulse duration of ~80 fs and a repetition rate of 250 kHz excites a THz photoconductive (PC) emitter (Tera-SED10, Laser Quantum) and the generated THz rays are collimated by an off-axis parabolic mirror onto the kirigami structures at normal incidence. The spot size of the THz beam is controlled by an iris diaphragm to a diameter of ~2 cm to ensure we measure only THz waves passing through the kirigami modulator. The transmitted beam is focused by another set of parabolic mirrors and detected by a 1 mm thick (110)-oriented ZnTe crystal with the method of electro-optic (EO) sampling (*32*).

The following method utilizing two linear polarizers was used to determine the orientation and ellipticity of arbitrarily polarized THz waves (*6,7*). Two THz wire grid polarizers (G50 × 20-L, Microtech Instruments, Inc.) with an extinction ratio of $10^3$-$10^4$ in the spectral range 0.1-3 THz were used in the configuration shown in Fig. S6A. The THz fields generated by the PC emitter were measured to have a high degree of linear polarization with an ellipticity angle below 0.3° (shown as the dash-dot lines in Fig. S8 to Fig. S10), which is negligible compared to the ellipticity induced by the chiral kirigami modulator; we confirmed that use of a linear polarizer immediately after the emitter made no further improvement in linearity. The emitter was fixed at an orientation such that the generated THz polarization was horizontal to the optical table (defined here as the *x* axis). The first polarizer (P1) was placed in front of the ZnTe crystal and its transmission direction (perpendicular to the wire grid orientation) was fixed vertical to the optical table (defined as the *y* axis). The ZnTe crystal and the sampling pulses were also oriented to give the maximum electro-optic sensitivity along *y* direction. The second polarizer (P2) was placed between the sample and the first polarizer and was rotated to different orientations to determine the complete polarization state of the transmitted field.

When the P2 transmission axis is along the *y* direction (defined as 0°), it is aligned with P1 and the y-component of the transmitted waves through sample $E_y(t)$ is measured. The x-component $E_x(t)$ is measured by rotating the orientation of P2 to +45° and -45° and calculated by the subtraction of the two. Since any arbitrary electric field can be decomposed into two perpendicular components, polarization states such as ellipticity and polarization rotation angle can be fully determined with three measurements. The electric field from the PC emitter without samples was also measured using the same method for calculating the reference transmission coefficients and labeled as "emitter" in Fig. S8 to Fig. S10.

<u>Definition of sample mounting orientations and abbreviations for sample configurations</u>

Since our kirigami pattern does not have C4 symmetry, measurements must be performed for two perpendicular polarizations, *i.e*. horizontally and vertically polarized THz waves, incident on the kirigami to fully characterize the kirigami sheet's in-plane optical properties, especially for circular dichroism. This was accomplished by rotating the kirigami sheet by 90° instead of rotating the THz emitter, which would have required elaborate rotations of two polarizers as well as the ZnTe crystal and the sampling beam. The kirigami modulator was attached to a rotation mount (RSP1, Thorlabs), so the transmitted waves can be measured in both horizontal and vertical orientations (simple rotation by 90°). Here, the horizontal and vertical mounting orientations are defined as follows: (1) horizontal - stretching direction is along with x direction as indicated in Fig. S6B, (2) vertical - stretching direction is along with y direction in Fig. S6C.

There are four possible configurations for the measurements: the kirigami modulator may be designed for either right- or left-handedness, and the modulators may be mounted horizontally or vertically relative to the input linear polarization. The abbreviation used in this Supporting materials section is as follows: "*HL*" for horizontally mounted left-handed kirigami modulator, "*HR*" for horizontally right-handed, "*VL*" for vertically left-handed, and "*VR*" for vertically right-handed.

<u>Calculations of transmittance, polarization state and THz circular dichroism (TCD)</u>

As mentioned above, the *x*-component of the electric field can be calculated by

$$E_x(t) = E_{+45^\circ}(t) - E_{-45^\circ}(t) \tag{1}$$

where $E_{+45^\circ}(t)$ and $E_{-45^\circ}(t)$ are the time-domain electric field measurements when the transmission orientation of polarizer P2 are at +45° and -45° relative to that of the polarizer P1, respectively.

The electric field signals are measured in the time domain and the complex frequency-domain electric field spectra are obtained using fast Fourier transform (FFT)

$$\tilde{E}_x = \tilde{E}_x(\omega) = FFT\{E_x(t)\}$$
$$\tilde{E}_y = \tilde{E}_y(\omega) = FFT\{E_y(t)\} \tag{2}$$

The Jones transfer matrix of a sample can be defined as

$$T = \begin{pmatrix} t_{xx} & t_{yx} \\ t_{xy} & t_{yy} \end{pmatrix} \tag{3}$$

where the first subscript letter indicates the incident polarization direction and the second subscript indicates the output direction for detection; the electric field vector of the transmitted THz wave through the sample $\tilde{E}_s$ is related to the incident electric field $\tilde{E}_{in}$ by

$$\tilde{E}_s = T\tilde{E}_{in} \tag{4}$$

The electric field $\tilde{E}_{in}$ incident on the sample is the reference electric field generated by the PC emitter measured without a sample $\tilde{E}_{ref}$ and in our measurement, it is always along the *x* direction. When the sample is mounted horizontally, electric field and transmission coefficients have relation as follow:

$$\begin{pmatrix} \tilde{E}_{sx}^h \\ \tilde{E}_{sy}^h \end{pmatrix} = \begin{pmatrix} t_{xx} & t_{yx} \\ t_{xy} & t_{yy} \end{pmatrix} \begin{pmatrix} \tilde{E}_{ref} \\ 0 \end{pmatrix} \tag{5}$$

Here, superscripts indicate the abbreviation of mounting orientation. Also, the two transmission coefficients of the sample at horizontal position can be calculated by

$$t_{xx} = \tilde{E}_{sx}^h \,/\, \tilde{E}_{ref}$$
$$t_{xy} = \tilde{E}_{sy}^h \,/\, \tilde{E}_{ref} \tag{6}$$

After rotating the sample by 90° which is defined as vertical orientation, the measured transmitted signals are related to the reference signal as

$$\tilde{E}_s^v = R(90°)TR(-90°)\tilde{E}_{ref} \tag{7}$$

where $R(\theta)$ is the rotation matrix with rotation angle $\theta$

$$R(\theta) = \begin{pmatrix} cos(\theta) & -sin(\theta) \\ sin(\theta) & cos(\theta) \end{pmatrix} \tag{8}$$

Therefore, the measured transmitted signals of the sample for vertical orientation are

$$\begin{pmatrix} \tilde{E}_{sx}^{v} \\ \tilde{E}_{sy}^{v} \end{pmatrix} = \begin{pmatrix} 0 & -1 \\ 1 & 0 \end{pmatrix} \begin{pmatrix} t_{xx} & t_{yx} \\ t_{xy} & t_{yy} \end{pmatrix} \begin{pmatrix} 0 & 1 \\ -1 & 0 \end{pmatrix} \begin{pmatrix} \tilde{E}_{ref} \\ 0 \end{pmatrix}$$
$$= \begin{pmatrix} t_{yy} & -t_{xy} \\ -t_{yx} & t_{xx} \end{pmatrix} \begin{pmatrix} \tilde{E}_{ref} \\ 0 \end{pmatrix} \tag{9}$$

and the two transmission coefficients of the sample at vertical orientation can be calculated by

$$t_{yy} = \tilde{E}_{sx}^{v} \;/\; \tilde{E}_{ref}$$
$$t_{yx} = -\,\tilde{E}_{sy}^{v} \;/\; \tilde{E}_{ref} \tag{10}$$

Since the polarization of the incident THz beam is linear and horizontal, the sample-induced polarization rotation angle $\theta$ and ellipticity angle $\eta$ (*33*) can be calculated directly by the measured THz spectra of $\tilde{E}_s = \begin{pmatrix} \tilde{E}_x \\ \tilde{E}_y \end{pmatrix}$ using Stokes parameters (*33*), and the same equations can be applied for both the horizontal and vertical orientations of mounting. The four Stokes parameters are defined as

$$S_0 = \tilde{E}_x\tilde{E}_x^* + \tilde{E}_y\tilde{E}_y^*$$
$$S_1 = \tilde{E}_x\tilde{E}_x^* - \tilde{E}_y\tilde{E}_y^*$$
$$S_2 = \tilde{E}_x\tilde{E}_y^* + \tilde{E}_y\tilde{E}_x^*$$
$$S_3 = i(\tilde{E}_x\tilde{E}_y^* - \tilde{E}_y\tilde{E}_x^*) \tag{11}$$

Since THz-TDS measures the electric field directly, three measurements (one for $\tilde{E}_y$and two for $\tilde{E}_x$) determine the four Stokes parameters and thus the polarization state.

The polarization rotation angle $\theta$ relative to the horizontal direction and the ellipticity $\eta$ can be calculated using Stokes parameters as follow:

$$\theta = \frac{1}{2} tan^{-1}\left(\frac{S_2}{S_1}\right), \qquad -\frac{\pi}{2} \leq \theta \leq \frac{\pi}{2}$$
$$\eta = \frac{1}{2} sin^{-1}\left(\frac{S_3}{S_0}\right), \qquad -\frac{\pi}{4} \leq \eta \leq \frac{\pi}{4} \tag{12}$$

Additional care should be taken for the rotation angle $\theta$, because mathematically the range of the inverse tangent function $tan^{-1}(x)$ is $\left[-\frac{\pi}{2},\frac{\pi}{2}\right]$ and correspondingly the range of $\theta$ would be $\left[-\frac{\pi}{4},\frac{\pi}{4}\right]$. In optics, however, the rotation angle $\theta$ is within the range of $\left[-\frac{\pi}{2},\frac{\pi}{2}\right]$. This can be easily illustrated via the Poincaré sphere in which $2\theta$ covers a whole circle, *i.e.* from $-\pi$ to $\pi$. The three perpendicular axes on the Poincaré sphere can be represented by the three Stokes parameters $S_1$, $S_2$ and $S_3$, so the following conditions are used to convert the mathematical inverse tangent function given by the numerical computing software (MATLAB for this paper) to the actual optical rotation angle $\theta$

$$\begin{aligned}
&\text{if } S_1 \geq 0: \quad \theta = \frac{1}{2}tan^{-1}\left(\frac{S_2}{S_1}\right), \qquad -\frac{\pi}{4} \leq \theta \leq \frac{\pi}{4} \\
&if\ S_1 < 0 \text{ and } S_2 \geq 0: \theta = \frac{1}{2}tan^{-1}\left(\frac{S_2}{S_1}\right) + \frac{\pi}{2}, \qquad \frac{\pi}{4} < \theta \leq \frac{\pi}{2} \\
&if\ S_1 < 0\ and\ S_2 < 0: \theta = \frac{1}{2}tan^{-1}\left(\frac{S_2}{S_1}\right) - \frac{\pi}{2}, \qquad -\frac{\pi}{2} \leq \theta < -\frac{\pi}{4}
\end{aligned} \tag{13}$$

Alternatively, it can be directly calculated by using the four-quadrant inverse tangent function, especially for MATLAB.

$$\theta = \frac{1}{2}atan2(S_2,\ S_1), \qquad -\frac{\pi}{2} \leq \theta \leq \frac{\pi}{2} \tag{14}$$

The transmitted electric field through a kirigami sample for a circularly polarized incident beam can be inferred using the Jones matrix elements measured from linearly polarized incident fields. For a normalized right circularly polarized (RCP) incident beam (*34*)

$$\tilde{E}_{RCP}^{in} = \frac{1}{\sqrt{2}}\begin{pmatrix} 1 \\ i \end{pmatrix} \tag{15}$$

the electric field of the transmitted wave is

$$\tilde{E}_{RCP}^{out} = \begin{pmatrix} t_{xx} & t_{yx} \\ t_{xy} & t_{yy} \end{pmatrix} \frac{1}{\sqrt{2}}\begin{pmatrix} 1 \\ i \end{pmatrix} = \frac{1}{\sqrt{2}}\begin{pmatrix} t_{xx} + it_{yx} \\ t_{xy} + it_{yy} \end{pmatrix} \tag{16}$$

and the magnitude of this complex electric field vector is

$$E_R = \frac{1}{\sqrt{2}}\sqrt{\left|t_{xx} + it_{yx}\right|^2 + \left|t_{xy} + it_{yy}\right|^2} \tag{17}$$

where | | is the absolute value of a complex number.
Similarly, for a normalized left circularly polarized (LCP) incident beam (*34*)

$$\tilde{E}^{in}_{LCP} = \frac{1}{\sqrt{2}}\begin{pmatrix} 1 \\ -i \end{pmatrix} \tag{18}$$

the electric field of the corresponding transmitted wave is

$$\tilde{E}^{out}_{LCP} = \begin{pmatrix} t_{xx} & t_{yx} \\ t_{xy} & t_{yy} \end{pmatrix}\frac{1}{\sqrt{2}}\begin{pmatrix} 1 \\ -i \end{pmatrix} = \frac{1}{\sqrt{2}}\begin{pmatrix} t_{xx} - it_{yx} \\ t_{xy} - it_{yy} \end{pmatrix} \tag{19}$$

and the magnitude of this complex electric field vector is

$$E_L = \frac{1}{\sqrt{2}}\sqrt{\left|t_{xx} - it_{yx}\right|^2 + \left|t_{xy} - it_{yy}\right|^2} \tag{20}$$

The terahertz circular dichroism (TCD) is a commonly used quantity for characterizing the optical activity of chiral materials (*35*). It is related to the relative transmission (or absorption) difference between RCP and LCP incident waves, and can be defined and quantified by (*35*)

$$TCD = tan^{-1}\left(\frac{E_R - E_L}{E_R + E_L}\right) \tag{21}$$

where $E_R$ and $E_L$ are the magnitudes of the transmitted waves of RCP and LCP incident beams given by Eq. (17) and Eq. (20), respectively.

<u>An example of raw THz-TDS data for kirigami sheets</u>

As an example of the raw data measured directly from THz-TDS with the two-linear polarization setup, Fig. S7A shows the three time-domain THz electric fields (i.e. with the polarizer P2 rotated to +45°, -45° or 0° position) of a horizontally mounted left-handed (*HL*) kirigami sample with gold herringbone pattern at $\varphi$ = 30° and stretched with ε = 22.5%. Fig. S7B are the electric fields for the same sample under the same measurement conditions but rotated to vertical (*VL*). Fig. S7C and D show zoomed views near zero time delay, and the signals of the 0° component and the appearance of multi-cycle waves compared to single-cycle input pulse (especially for the 0° component) are clear evidences of polarization rotation and THz resonances of chiral kirigami structures. These raw data were then processed using the equations presented in the previous section to calculate the electric field transmittances, polarization angles and circular dichroism.

Experimental data of transmittance, polarization state and TCD of single kirigami modulator

The detailed data for kirigami modulator with three different slant angles $\varphi$ are presented in Fig. S8 to Fig. S11. These include the spectra of the magnitudes of the transmission coefficients ($t_{xx}$, $t_{xy}$, $t_{yx}$, $t_{yy}$), the polarization rotation ($\theta$) and ellipticity angles ($\eta$) for the different slant angles ($\varphi$) for both horizontal and vertical mounts, and the spectra of the resulting circular dichroism angles. In addition, kirigami structures with narrower gold wire widths and spacings (2 μm wire width and 2 μm spacing) but otherwise the same design parameters are also measured to explore the origin of resonance. Unless stated otherwise, all other data presented in the main text and supplementary part were measured with samples with gold strips with 5 μm wire width and 5 μm spacing.

The largest strain applied in these measurements was $\varepsilon = 22.5\%$. This value was chosen for the optimum performance as well as maintaining the function as modulators for the kirigami samples. It was measured that beyond this strain the polarization rotation and ellipticity saturated. With a much larger strain, the signal actually decreased because the Au strip surfaces became parallel to the THz beam propagation direction and the effective interaction area became smaller. In addition, a very large strain would deform the sample out of the elastic range and deteriorate the function of these samples as modulators.

Measurements and modeling of kirigami 3D geometries

The 3D topography of the kirigami sheets under various strains were measured using a laser confocal microscope (OLS 4000 LEXT, Olympus). The kirigami geometries were reconstructed using 3D graphic software packages (Rhino 5, Robert McNeel & Associates and 3D Max 2017, Autodesk) based on the experimentally acquired images from the confocal microscopy. Three projected images captured at different view angles were used to clearly define and reconstruct the radius, angles, shapes and boundaries. Two 3D structures of right-handed (*R*) kirigami sheets with 45° slanted Au strips were made as shown in Fig. S5A and B. Kirigami sheets were flipped to make reversed handedness, left-handed (*L*) structures. In addition, an achiral structure with a 0° wire slanted angle was also modeled using the same procedure as shown in Fig. S5C.

TCD simulation and surface current norm distributions

TCD simulations and surface current norm distributions of illuminated kirigami modulators were numerically investigated with finite element simulation software (COMSOL Multiphysics 5.2a, COMSOL Inc.) using the reconstructed 3D models for the shape. Periodic boundary conditions were applied to the single unit cell of each model to simplify the simulation and to reduce the required computational power. It should be noted that this could lead to some simulation errors due to the infinite unit cells assumed by periodic boundary conditions. In the experiment, of course, there are a finite number of unit cells defined by diameter of the THz beam; this could be one of the reasons that the magnitudes of the simulation were slightly larger than the experimental data. Other possible reasons for this slight difference between experiment and simulation can be the frequency resolution limit of the experimental setup (*6*) and the errors generated during the modeling process and inherent limitations of COMSOL solvers. Apart from this slight difference, the simulated results matched well with the experimental data for the circular dichroism signs, peak positions, peak width and magnitudes.

Circuit model for calculating resonance frequency

The inductance of $n$ parallel conductors with identical dimensions can be calculated as

$$L = \sum_{i=1}^{n} L_i + \sum_{i=1}^{n} \sum_{j=1, j\neq i}^{n} M_{i,j} \tag{22}$$

where $L_i$ is the self-inductance and $M_{i,j}$ is the mutual inductance (*36*). Here we assume that each conductor carries the same current and the current is uniformly distributed over the entire cross section. To obtain a simple expression, we convert $n$ identical parallel conductors into one large single equivalent current sheet having total width of $\rho = nw + (n-1)s$, where $s$, $w$, $l$ are edge-to-edge spacing, width and length of the strips, respectively as described in reference (*36*). The result of sheet approximation is

$$L \approx \frac{\mu_0 n^2 l}{2\pi}\left[\ln\left(\frac{2}{\rho}\right) + 0.5 + \frac{\rho}{3} - \frac{\rho^2}{24}\right] \tag{23}$$

where $\mu_0$ is the permeability of free space and variable $\rho$ is the ratio of the width of the equivalent current sheet to its length. In addition, the self-capacitance of the Au strips is a combination of the capacitance between the gap capacitance in air and substrate. The total capacitance of the parallel conductors is given below:

$$C = l(n-1)\left(\frac{\varepsilon_0}{2}\frac{K(k')}{K(k)} + 2\varepsilon_0\varepsilon_r\frac{t}{s}\right) \quad and \quad k' = \sqrt{1-k^2} \tag{24}$$

where $\varepsilon_0$ is the permittivity of free space and $\varepsilon_r$ is the relative permittivity. $K(k)$ represents the elliptical integral of first order to calculate the effect of the fringing field. The modulus $k$ defined as $k = cos(\frac{\pi}{2}\frac{w}{s+2})$ is determined by the periodic geometry (*37*). The scaling of the optical response of parallel metal strips can be obtained by modeling the structure as an $LC$ circuit with the resonance frequency of $f_r = (2\pi\sqrt{LC})^{-1}$ (*27*). Roughly, inductance $L$ and capacitance $C$ scale linearly with the length of Au strips, $l$, and therefore $f_r$ become inversely proportional to $l$ as shown in Fig. 3d in the main text.

Experimental setup and data of polarization states for stacked kirigami modulators

Before measuring TCD using kirigami modulator, we studied polarization states for double stacked kirigami sheets. Kirigami chiroptical modulators with 37.5° slant angle ($\varphi$) were used to characterize and demonstrate the effects of stacking two kirigami layers on the polarization states. Each individual modulator was measured separately using the two-polarizer method described above, with results shown in Fig. S13. From these data, circular dichroism spectra for left- (*L*) and right- (*R*) handed samples were calculated with results shown in Fig. S14 using the same equation stated previously. Two kirigami modulators were then stacked along the collimated THz propagation (*z*-axis) direction, with a separation of ~2 cm and actuated by two independent piezo-controllers from 0 % to 18 % strains, as shown by the schematic of the experimental setup in Fig. S15. Four combinations of two samples (i.e. *VL* and *HL*, *VR* and *HL*, *VL* and *HR*, *VR* and *HR*) were measured and the total effects on the rotation and ellipticity of the polarization are shown in Fig. S16 to Fig. S19. These four combinations were chosen to show the *enhancement* of optical activity by stacking kirigami with the *same* chirality and *compensation* by stacking kirigami with *opposite* chirality. This measurement also validates the use of kirigami modulators for THz circular dichroism spectroscopy. It turns out that if the mounting orientations for the two kirigami modulators are perpendicular i.e. the first is oriented vertically (*V*) and the second horizontally (*H*), the birefringence cancels. This is an important requirement for the combinations of modulators with opposite chirality to compensate *both* optical activity and birefringence. This information may be useful for future application. Also, for each combination there were 25 modulations achieved by 5 different strains applied to each kirigami independently as shown in Fig. S16-S19.

The results of Fig. S17 (*VR* and *HL*) and Fig. S18 (*VL* and *HR*) indicate that chirality-switchable modulator can be achieved by stacking kirigami modulators with opposite handedness. Moreover, it is only when the strains applied to the two layers are the same that the output polarization state is the same as the input, i.e. zero polarization rotation and zero ellipticity. The small non-zero values at these strains in the experimental results came from the imperfect matching and alignment between the two samples and can be improved by more careful control of samples or by additional calibration methods. In general, the experimental results match the theory very well, and this has the potential to be developed as a modulator for other applications such as secure THz communication and handedness-switchable devices.

The results of stacking kirigami modulators with the same chirality (Fig. S16 (*VL* and *HL*) and Fig. S19 (*VR* and *HR*)) indicate that adding polarization rotation and ellipticity can be achieved. The magnitudes are larger than for a single kirigami for all strain conditions except $\varepsilon$ = 0 % (Fig. S13). This shows the possibility that ideal 90° rotation angle and 45° ellipticity could be achieved by stacking more kirigami layers or using kirigami with parameters accurately designed for specific frequency.

TCD of kirigami sample using kirigami modulator

To demonstrate TCD application, we tested using double stacked kirigami ($\varphi$ of 37.5°) configuration (Fig. S15) - one kirigami as a chiroptical modulator (1st kirigami) and the other (2nd kirigami) as a tunable sample. Dynamic modulations of the two chiral kirigami were applied independently which translated into individual manipulations of ellipticity of the input beam (from linear to elliptical) and the chiroptical activity of the sample (from achiral to chiral). The first kirigami modulator generates left- or right- elliptical/circular polarization beams similar to a photoelastic modulator (PEM) in a conventional CD spectrometer (*35*), and the second kirigami can be considered as the sample to be probed.

To avoid confusion with the previously defined TCD angle in Eq. (21), which is calculated using ideal circular polarization deduced from the experimental linear polarization measurements, the TCD directly obtained by kirigami modulation is measured directly from the elliptically polarized beam generated by first kirigami. Since the elliptically polarized beam generated by a first kirigami is frequency dependent, the TCD angle is defined similarly to Eq. (21) above, but slightly modified to

$$TCD\ by\ kirigami\ modulator\ = tan^{-1}\left(\frac{E_{LK} - E_{RK}}{E_{LK} + E_{RK}}\right) = tan^{-1}\left(\frac{{I_{LK}}^{1/2} - {I_{RK}}^{1/2}}{{I_{LK}}^{1/2} + {I_{RK}}^{1/2}}\right) \qquad (25)$$

where $E_{LK}$ and $E_{RK}$ are the electric field magnitudes of the transmitted waves through the kirigami sample (the second kirigami in this case) of the elliptically polarized beam generated by the first kirigami sheet, which may be left-handed kirigami (*L*) or right-handed kirigami (*R*), respectively. $I_{LK}$ and $I_{RK}$ are the corresponding transmittance-intensities.

The transmittance-intensities were obtained and normalized to that of ε = 0%, which is with no strain applied to the kirigami modulators. The transmittance-intensities of ε = 0% is used as base to eliminate the inherent difference caused by any slight mismatch of *L*- and *R*- elliptically polarized beams. Since the modulation of the two chiral kirigami structures can be manipulated independently, we can test the measurement of TCD by manipulating both the ellipticity of the input polarization from linear to 20° elliptical and the chiroptical activity of sample to be probed from achiral to chiral. Fig. S20 shows the experimental data of TCD spectra of *L*- and *R*- kirigami samples with 5 strain conditions, for 5 different input elliptical polarizations. When the strain applied on the second kirigami increased, the measured TCD also increased, just as for the results in single kirigami measurements (Fig. S14). The measured TCD increased as the ellipticity of the input beam became larger because the intensity difference between the incident left- and right-elliptically polarized beams is increased. The top row subfigures all have zero TCDs because they were normalized by themselves using the processing method mentioned above.

Validation of our method is performed by comparing this TCD measured by kirigami modulator (Fig. S14B) with TCD measured by standard two polarizers method (Fig. S14A). This

TCD measurement on sample kirigami sheets demonstrates the potential of this method for vibrational CD (VCD) measurements of chiral bio-molecular samples. It should also be emphasized that since only the intensities instead of electric field components are needed, compared to the method using two polarizers mentioned previously, this TCD spectroscopic method with chiral kirigami modulators provides more generalized applications beyond THz-TDS such as integration with conventional Fourier-transform infrared spectroscopy (FTIR) for vibrational CD (VCD) measurement and THz camera for real-time polarization resolved 2D images.

TCD of biological samples using kirigami modulator

To further demonstrate the application of kirigami chiroptical modulators for TCD measurements of real biological samples, we measured TCD spectra of the elytron of a June beetle, a petal of a dandelion flower, a leaf of a maple tree and a piece of pig fat. Due to the small size of some biological samples, a focused THz beam with ~500 μm spot size was used and the schematic of the experimental setup is shown in Fig. S21. The whole setup was enclosed in a box purged by extra-dry nitrogen and the relative humidity was maintained below 3% to minimize the water vapor absorption and to maximize the measurement sensitivity. The kirigami chiroptical modulators with 37.5° slant angle were used to generate left- and right-handed elliptically polarized beams. Fig. S22 shows the results: (d) the reference, with no biological sample in place, shows near zero TCD indicating the intensity transmissions of the two kirigami modulators were almost the same; (e) the sample of a maple leaf shows a very small chiroptical response with slightly noisier curves, which mainly could come from the lower signal-to-noise ratio caused by the THz absorption by the leaf; (f) a petal of a dandelion flower shows TCD signals between 0.3-0.8 THz with negative value and with a TCD that increases as the input ellipticity gets larger; (g) the sample of pig fat also shows negligible TCD. The measured absorption coefficient (Fig. S23) of a dandelion shows that the absorption frequency range matches closely the TCD spectrum. The observed absorption in this frequency (0.3~0.8 THz) is different from that of the other biosamples, which may indicate that THz-active chemical constituents are likely different. Although we cannot rule out the possibility that TCD arises from the physical micro-structure of a petal, consistent chiral motifs are not observed in scanning electron microscope images of the sample. Rather, chemical constituents of a dandelion, such as caffeic acid, chlorogenic acid, chicoric acid, chrysoeriol and β-carotene are likely to have a substantive contribution (*38*), noting that the spectral positions of the TCD bands (Fig.S22) and those of absorption (Fig.S23) overlap the several vibration modes of caffeic acid (0.58 THz for monomer and 0.67 THz for dimer) (*39*), chlorogenic acid (0.41, 0.53, 0.7 and 0.9 THz) (*39*) and β-carotene (broad band from 0.25 to 2 THz) (*40*). As we show here, characterizing the chirality of matter through kirigami TCD spectroscopy could be the starting point for further studying of biological microstructure as well as a variety of biomolecules such as proteins and nucleic acids. Here, the measured transmittance, $T = \frac{I_{sam}(\omega)}{I_{ref}(\omega)} = \frac{(E_{sam}(\omega))^2}{(E_{ref}(\omega))^2}$, is obtained from the THz transmittance through a sample attached to an aperture, $I_{sam}(\omega) = (E_{sam}(\omega))^2$, divided by the THz transmittance through the void aperture, $I_{ref}(\omega) = (E_{ref}(\omega))^2$ . The absorption coefficient (α) is calculated by

$$\alpha(\omega) = -\frac{2}{d_s}\ln(T) \tag{26}$$

where $d_s$ is the thickness of the sample (*41*).

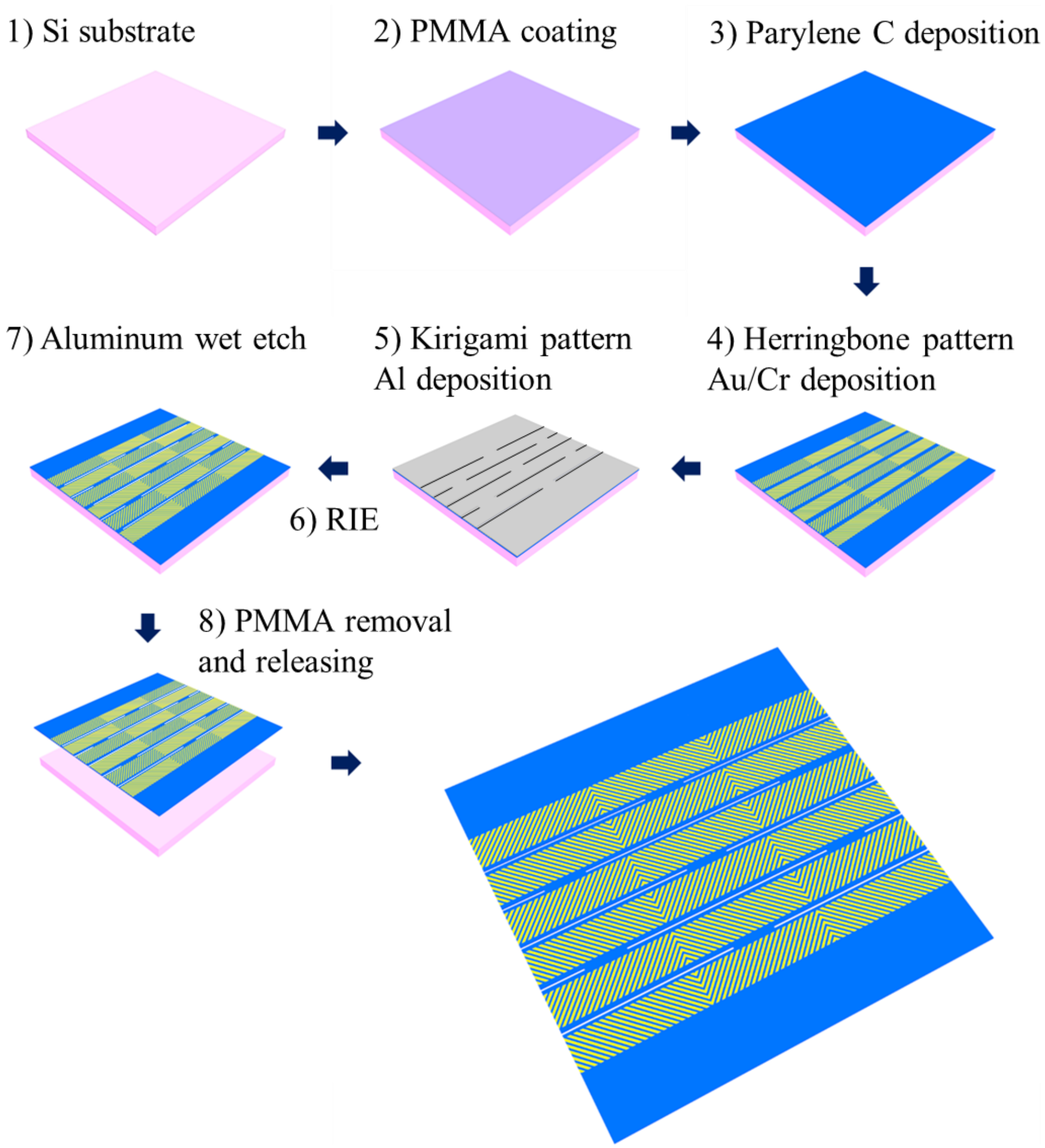


**Fig. S1. Fabrication of kirigami modulator.** Schematic illustration of the steps in fabrication processing. PMMA is applied as a sacrificial layer and subsequently ~6 μm thick Parylene C is deposited. Patterned Cr/Au layer is deposited by photolithography and electron beam evaporator. To introduce kirigami cut to Parylene C, additional Al layer is deposited for masking reactive ion etching (RIE). After RIE, the wafer is soaked in aluminum etchant solution and in acetone to release.

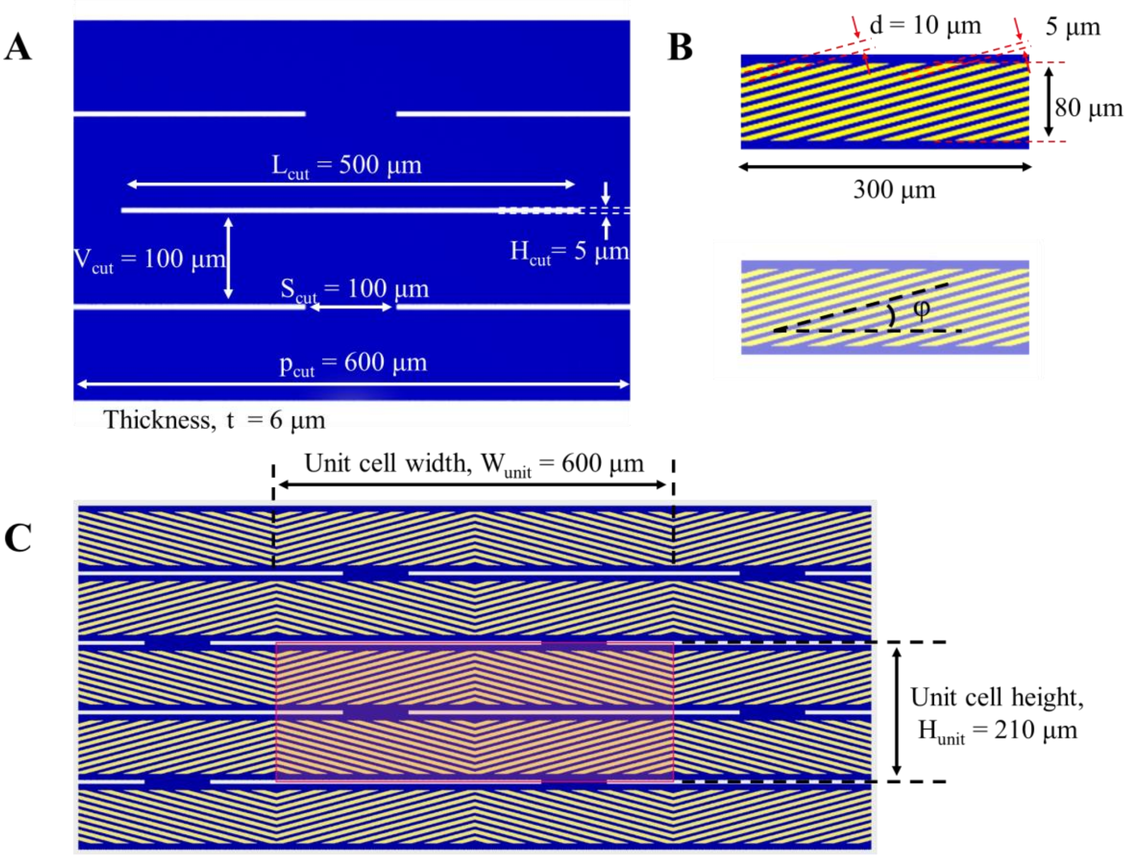


**Fig. S2. Detailed dimensions and definition of slant angle in chiral kirigami pattern.** (A) Top view of kirigami cut pattern. The length ($L_{cut}$) and the height ($H_{cut}$) of each individual cut are 500 μm and 5 μm, respectively. The horizontal ($S_{cut}$) and vertical ($V_{cut}$) spacing between cuts are set to 100 μm. (B) Detailed view of single unit of slanted Au strips. Width of each Au strip is set to 5 μm. Width and height of total domain of Au strips are 300 and 80 μm, respectively. Here, the slant angle ($\varphi$) is defined as angle between longitudinal direction of cut and Au strip as shown in Figure S2 below. (C) Top view image of aligned kirigami cut pattern and Au herringbone pattern. Red box indicates the unit cell of this double pattern. Width ($W_{unit}$) and height ($H_{unit}$) of unit cell are 600 and 210 μm, respectively.

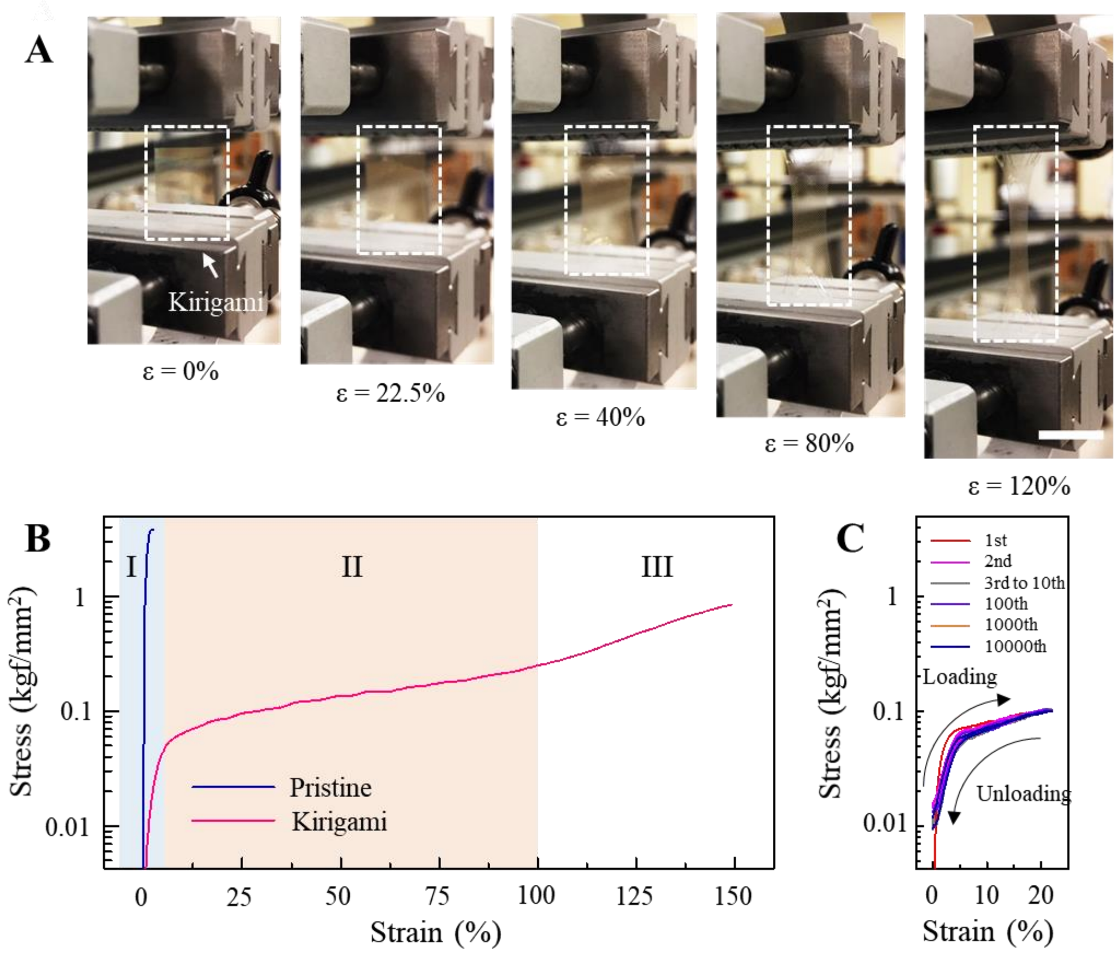


**Figure S3. Stretching and cycling properties of kirigami modulator** (A) Photo images of the kirigami at strain values of 0%, 22.5%, 40%, 80% and 120% (from left to right). (B) and (C) Stress-strain curves and their cycling properties of chiral kirigami modulator. Sections I (blue), II (pink) and III (white) indicate the regions of in-plane elastic deformation, out-of-plane elastic deformation and plastic deformation with pattern collapse, respectively. Scale bar in (A) is 2 cm.

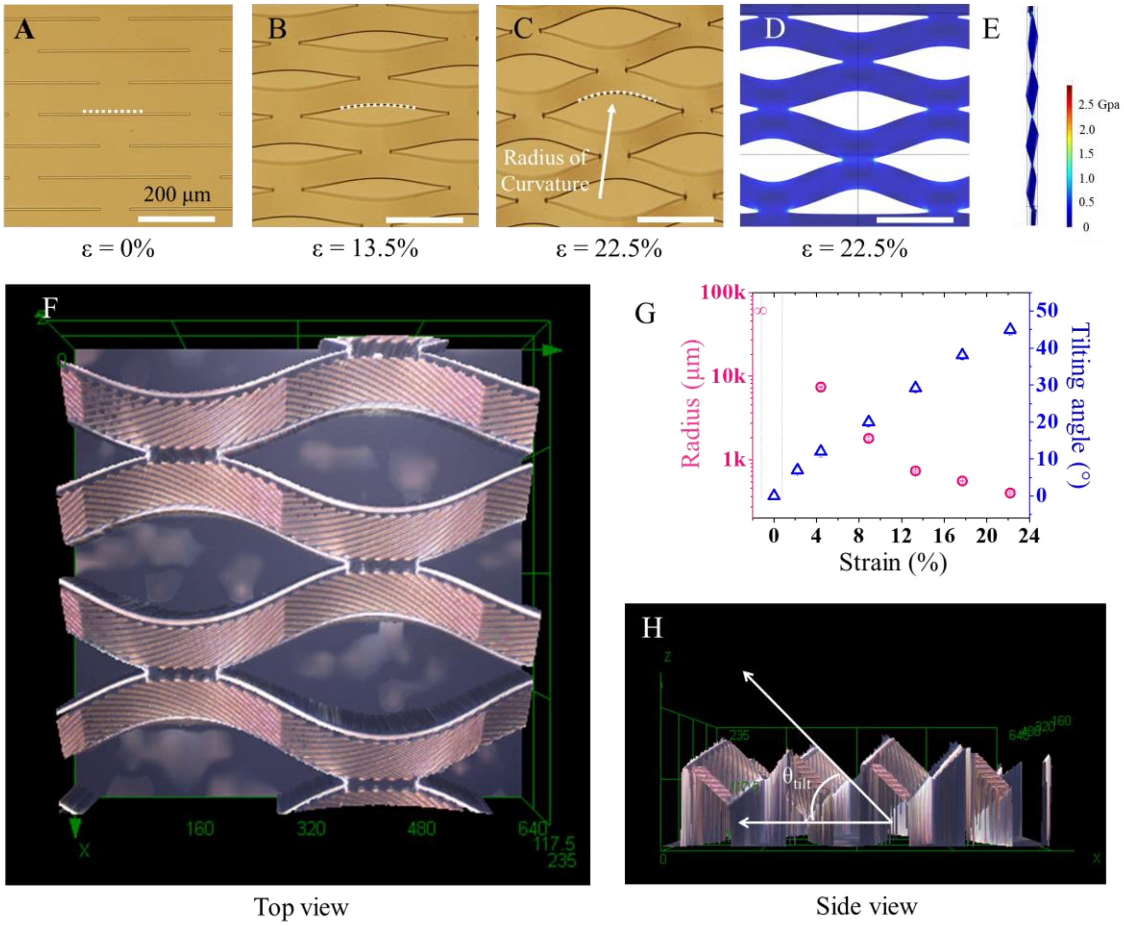


**Fig. S4. Structural evolution of kirigami modulator under tensile stress.** (A) to (C) show the optical microscope images of kirigami cut parylene at strain values of 0%, 13.5% and 22.5%, respectively. (D) and (E) show the top view and side view of stress distribution visualization in FEM, respectively. (F) and (H) show the top and side view kirigami at $\varepsilon$ = 22.5% strain captured by laser confocal microscopy, respectively. Here, the tilting angle ($\theta_{tilt}$) is defined as the angle between $x$ axis and the line parallel to the surface of the kirigami sheet as shown in (H). (G) shows the radius of the cut and tilting angle of the kirigami domain with respect to the strain (%). The radius of the cut edge was varied from almost infinity, i.e. flat line, to ~400 μm round while tilting angle changed from 0° to 45°.

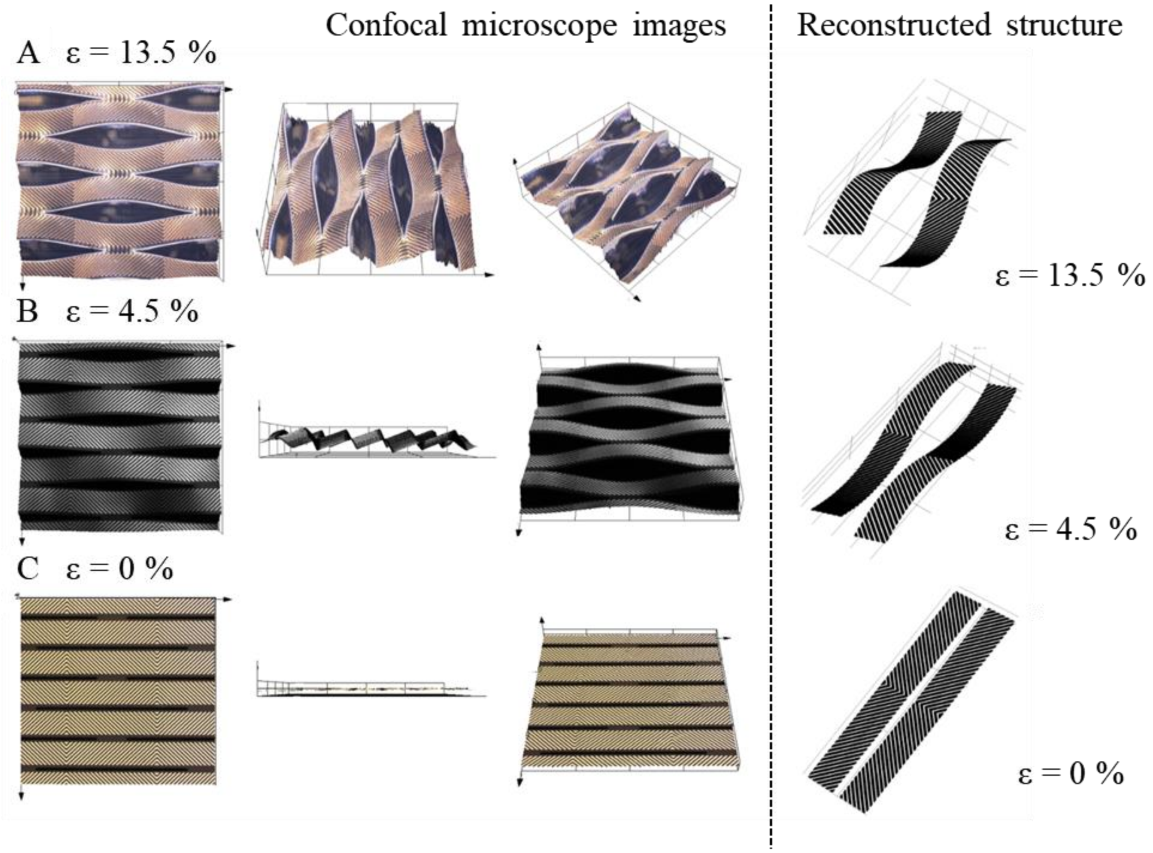


**Fig. S5. Confocal microscopy images and reconstructed 3D models of kirigami modulator under three different strains.** The three rows correspond to the same right-handed sample with a 45° wire slant angle under (A) 13.5%, (B) 4.5% and (C) 0% strains, respectively. The first three columns are the images of three different viewpoints from the confocal microscopy under a 20x objective. The last column are the reconstructed 3D models corresponding to each strain.

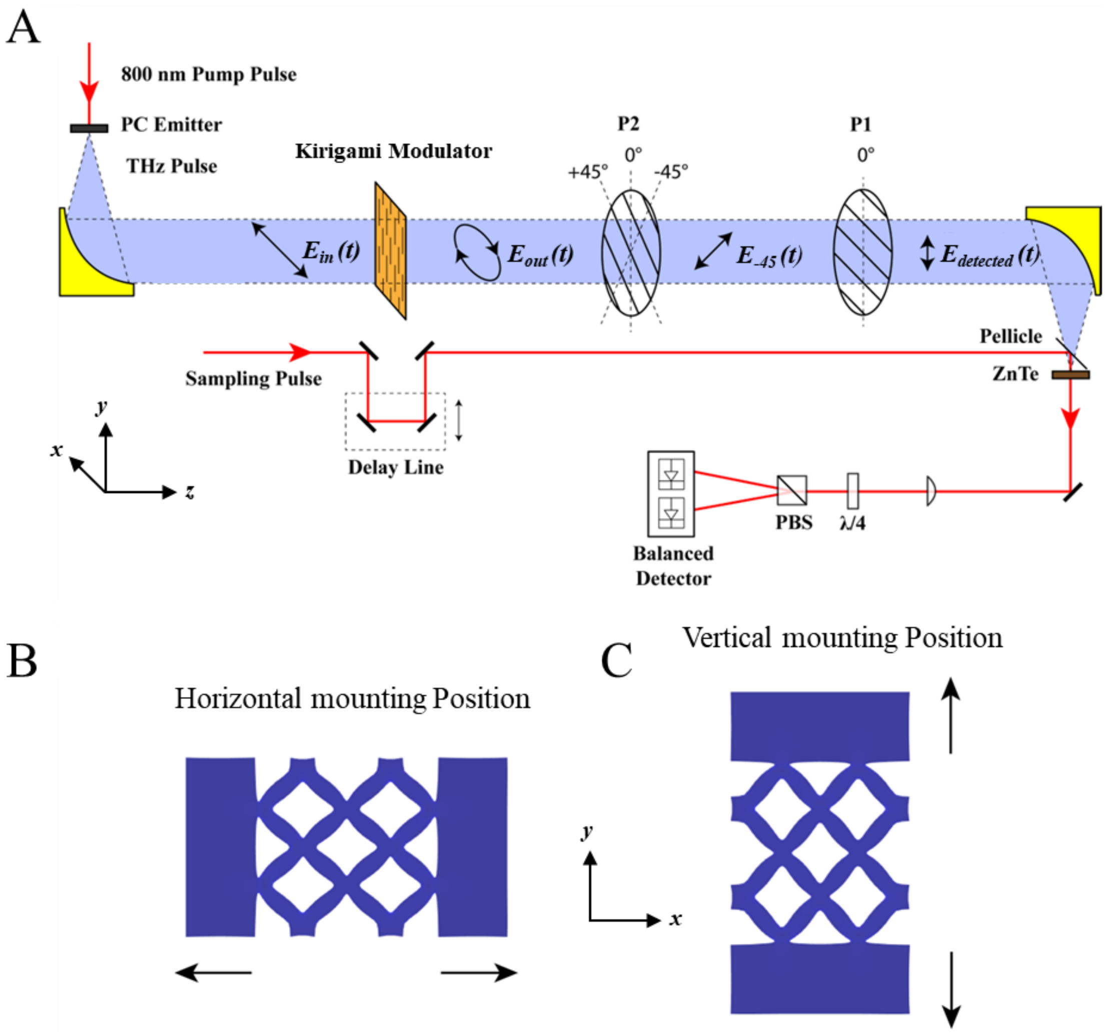


**Fig. S6. Schematics of the experimental setup and sample mounting positions for THz-TDS polarimetry measurement.** (A) Schematic of THz-TDS polarimetry measurement setup. The orientation of THz polarizer P1 is fixed at 0° to allow vertically polarized waves to transmit. The orientation of polarizer P2 is rotated to +45°, -45° or 0° for three polarization-selective measurements. This figure presents the orientation of P2 at -45° for example. (B) and (C) show the definitions of horizontal (H) and vertical (V) mounting positions. The thick black arrows indicate the stretching directions actuated by the piezo-controller horizontally for (B) and vertically for (C).

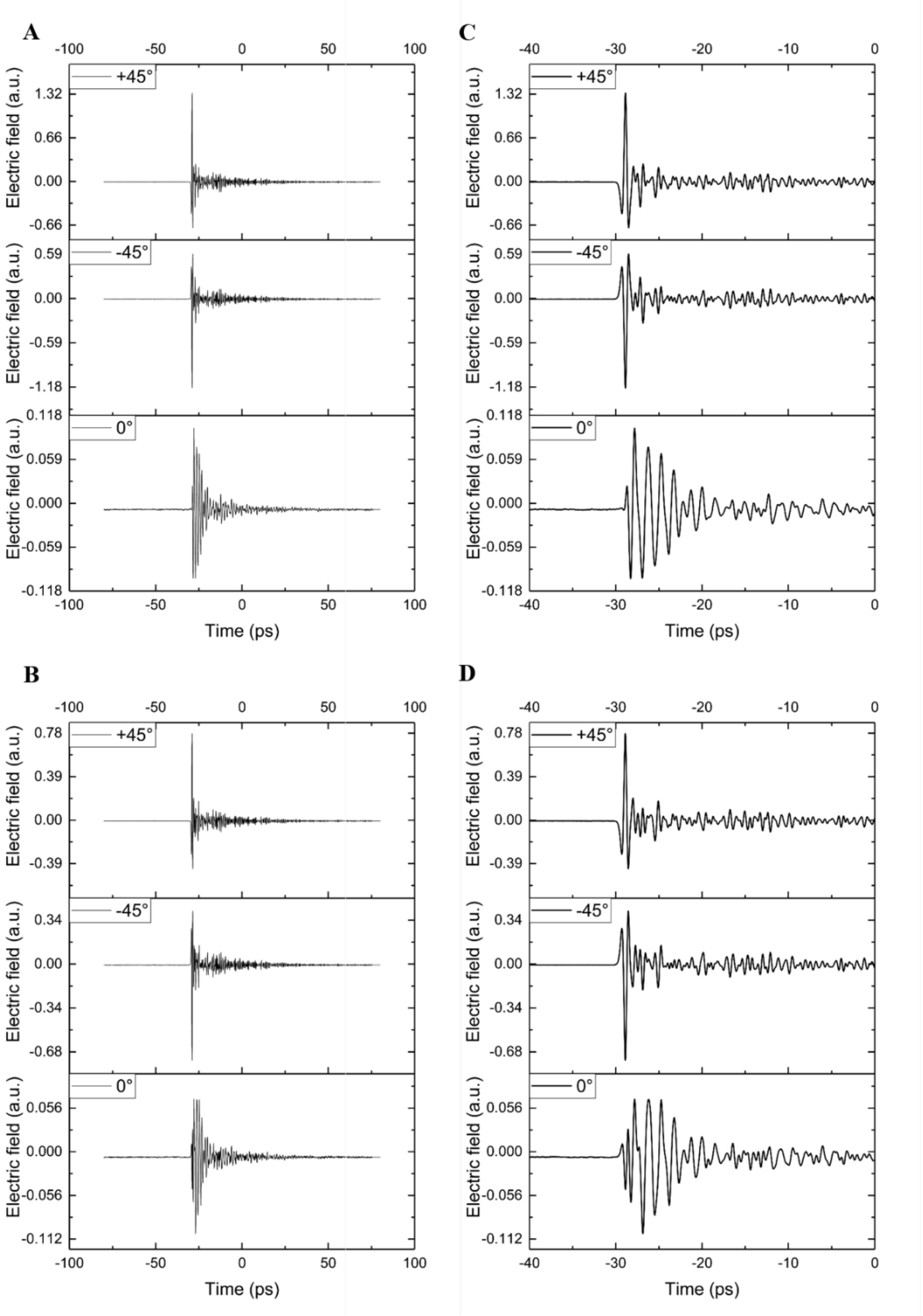


**Fig. S7. An example of raw THz-TDS data for three polarization measurements of a left-handed (*L*-) kirigami sample with 30° slant angle ($\varphi$ = 30°) and stretched with ε = 22.5%.** (A) Transmitted electric fields of horizontally and (B) vertically mounted sample over the whole scan range. (C) and (D) are zoomed views on the main peaks (near zero time delay) of (A) and (B), respectively.

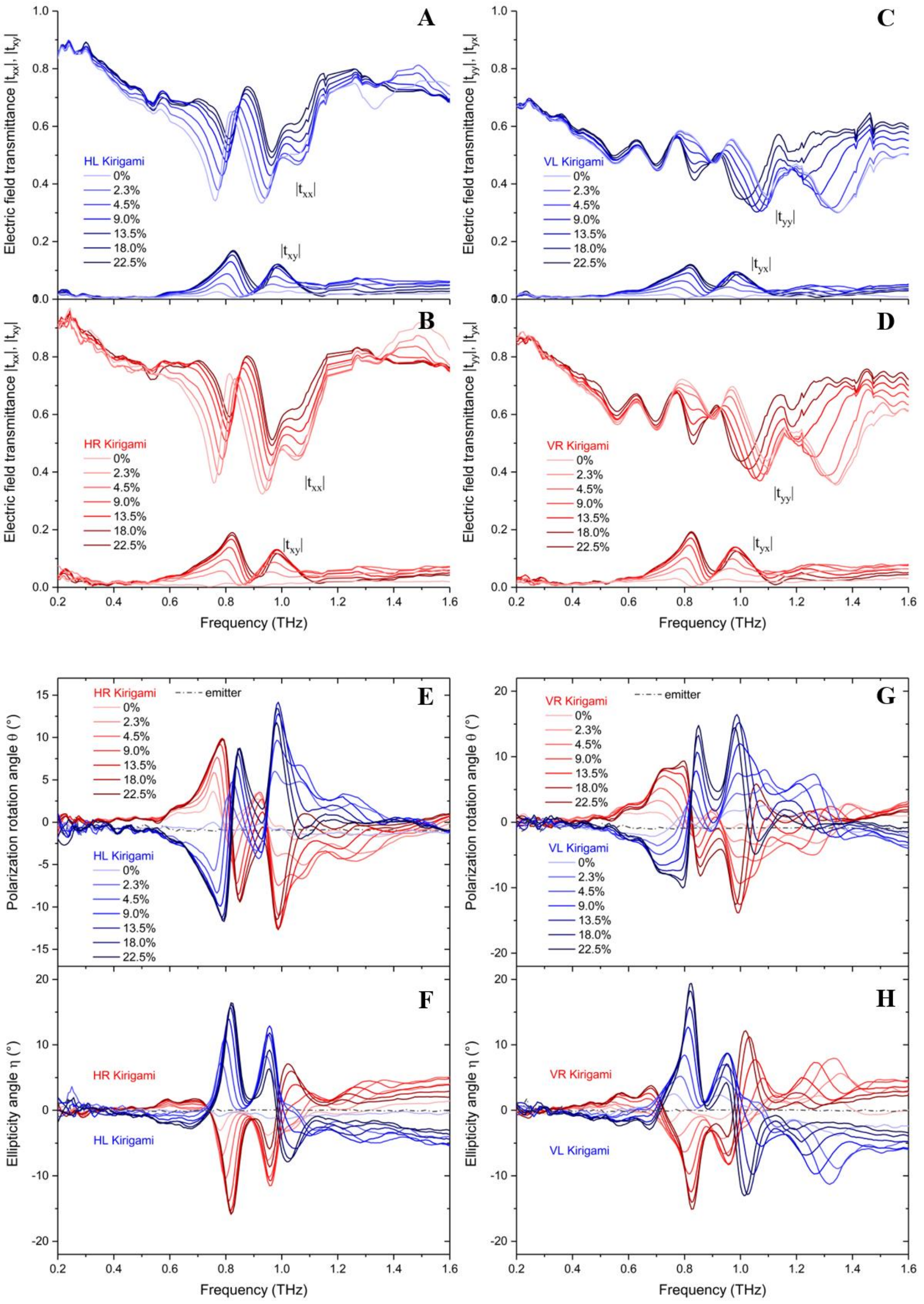


**Fig. S8. Experimental data for kirigami samples with wire slant angle of 45°.** (A) – (D) the magnitudes of four transmittance coefficients. (E) polarization rotation angle and (F) ellipticity angle induced by the samples mounted horizontally (H). (G) polarization rotation angle and (H) ellipticity angle induced by the samples mounted vertically (V). Blue and red curves are for left-handed (*L*) and right-handed (*R*) samples respectively. The strains applied are given in the legends and the same for all the subfigures.

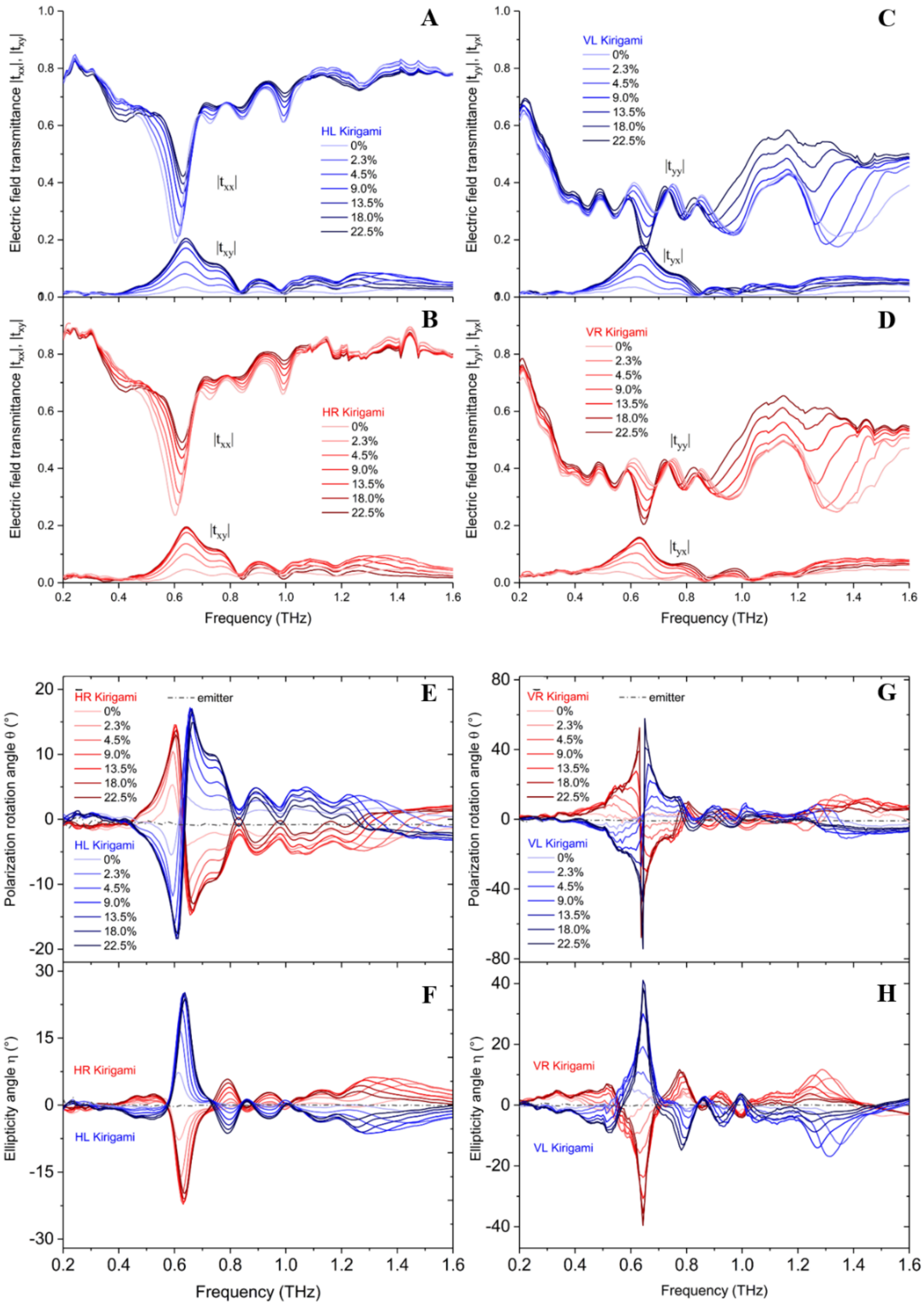


**Fig. S9. Experimental data for kirigami samples with wire slant angle of 30°.** (A) – (D) the magnitudes of four transmittance coefficients. (E) polarization rotation angle and (F) ellipticity angle induced by the samples mounted horizontally (H). (G) polarization rotation angle and (H) ellipticity angle induced by the samples mounted vertically (V). Blue and red curves are for left-handed (*L*) and right-handed (*R*) samples respectively. The strains applied are given in the legends and the same for all the subfigures.

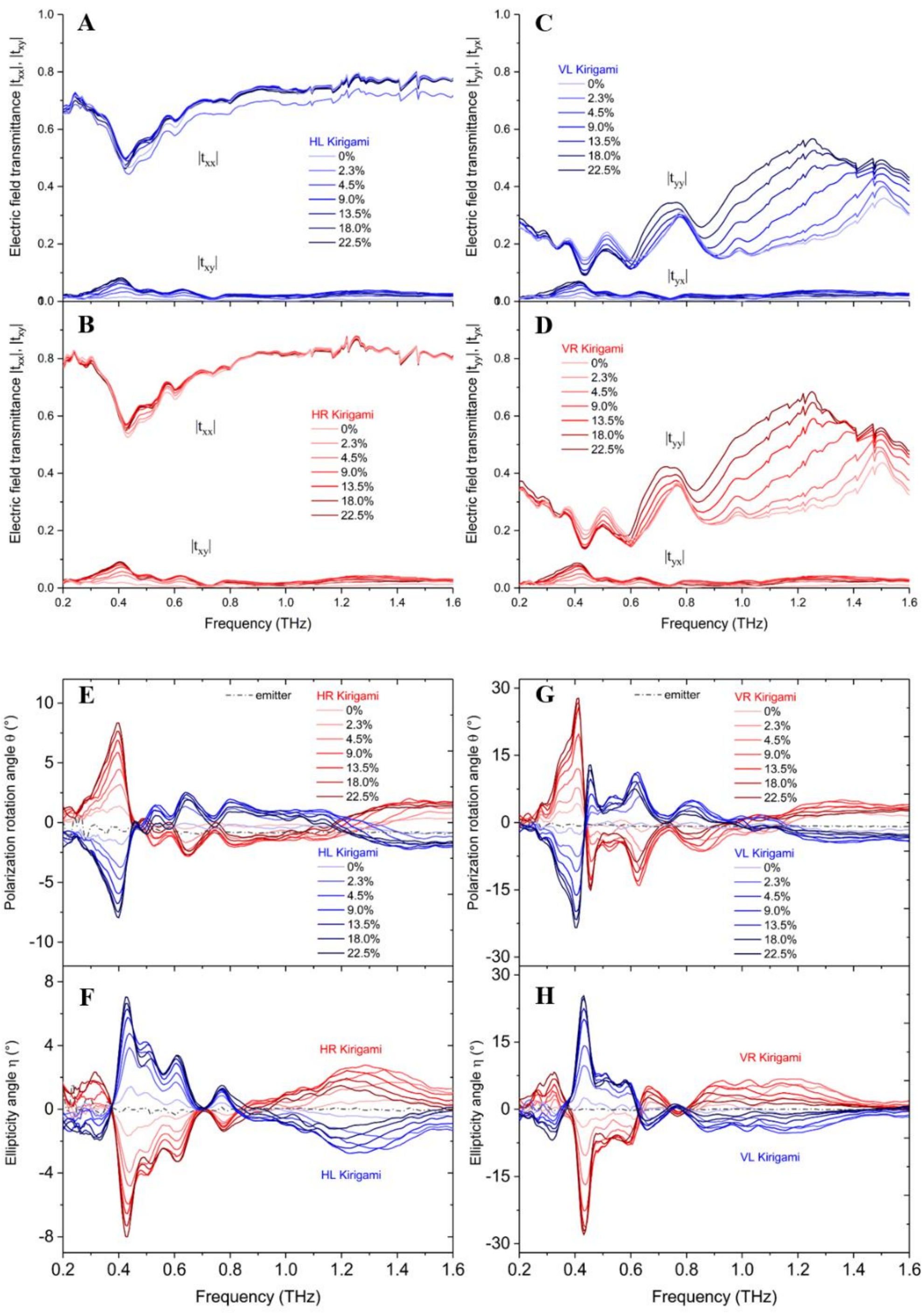


**Fig. S10. Experimental data for kirigami samples with wire slanted angle of 15°.** (A) – (D) the magnitudes of four transmittance coefficients. (E) polarization rotation angle and (F) ellipticity angle induced by the samples mounted horizontally (H). (G) polarization rotation angle and (H) ellipticity angle induced by the samples mounted vertically (V). Blue and red curves are for left-handed (*L*) and right-handed (*R*) samples respectively. The strains applied are given in the legends and the same for all the subfigures.

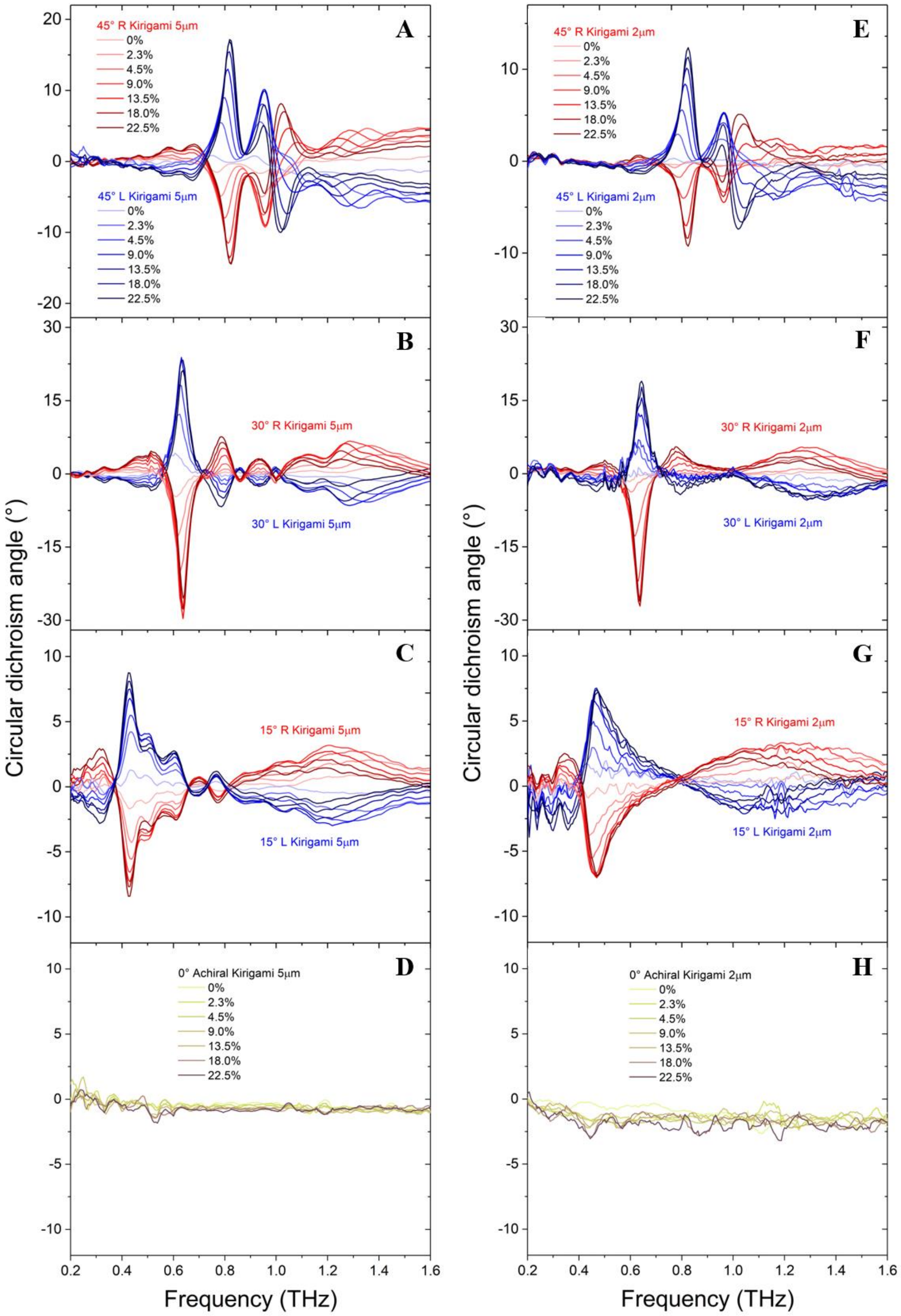


**Fig. S11. Comparison of circular dichroism spectra of kirigami samples with different gold wire widths and spacings.** The left (right) column corresponds to samples of 5 μm (2 μm) wire width and 5 μm (2 μm) spacing. The four rows correspond to samples with wire slant angles of 45° (A, E), 30° (B, F), 15° (C, G) and 0° (D, H), respectively. Blue and red curves are for left-handed (*L*) and right-handed (*R*) samples respectively and yellow curves are for achiral samples.

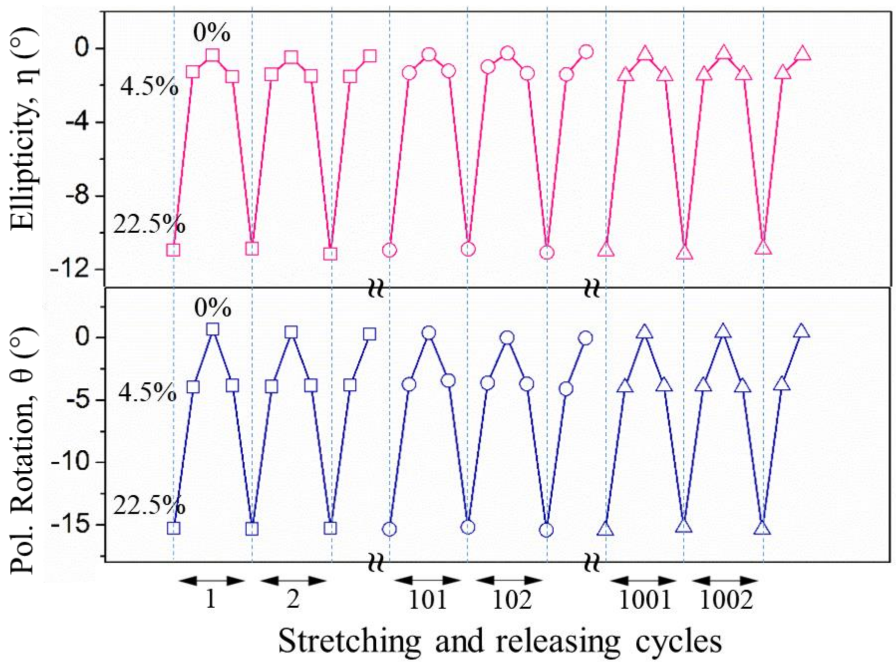


**Fig. S12. Results of polarization rotation and ellipticity angle of stretching and releasing cycles.** Cycling properties of polarization rotation and ellipticity angle. Both of $\theta$ and $\eta$ values are taken at 0.84 THz using kirigami modulator having $\varphi$ of 45°. Not only for mechanically but optically it maintains its values of polarization rotation and ellipticity angle even over 1000 cycles

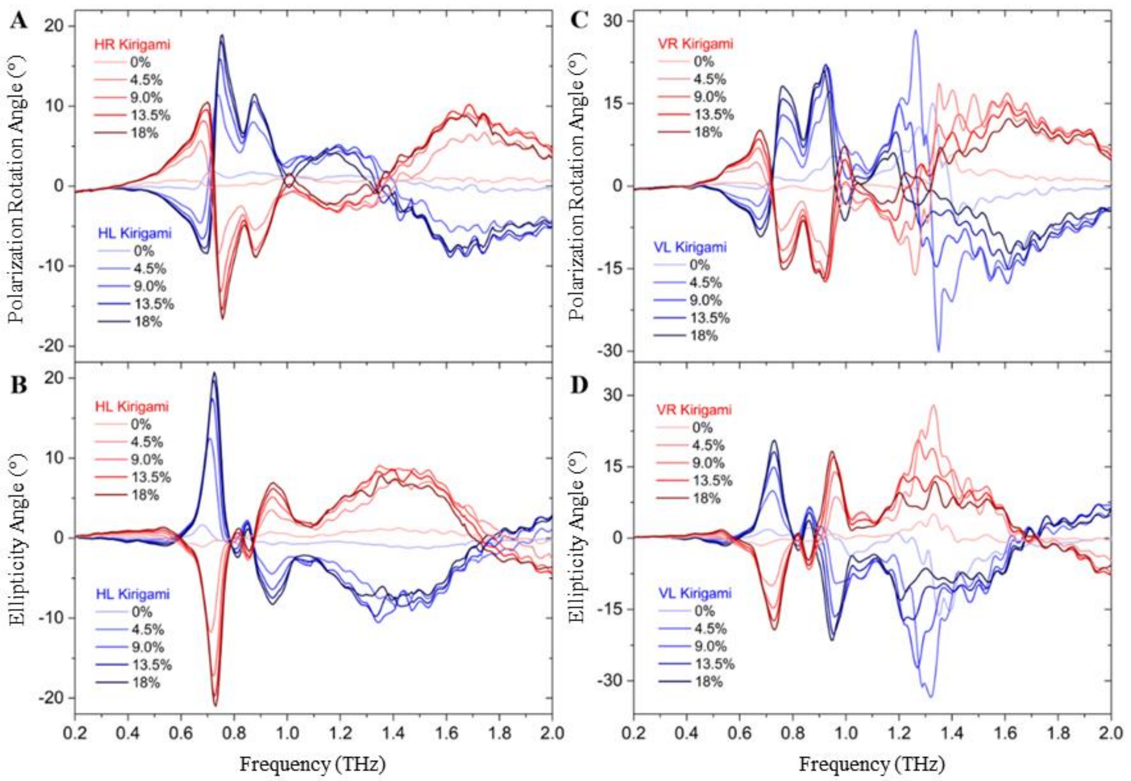


**Fig. S13. Experimental data for kirigami samples with slant angle of 37.5°.** (A) polarization rotation angle and (B) ellipticity angle induced by the samples mounted horizontally (H). (C) polarization rotation angle and (D) ellipticity angle induced by the samples mounted vertically (V). Blue and red curves are for left-handed (*L*) and right-handed (*R*) samples respectively. The strains applied are given in the legends.

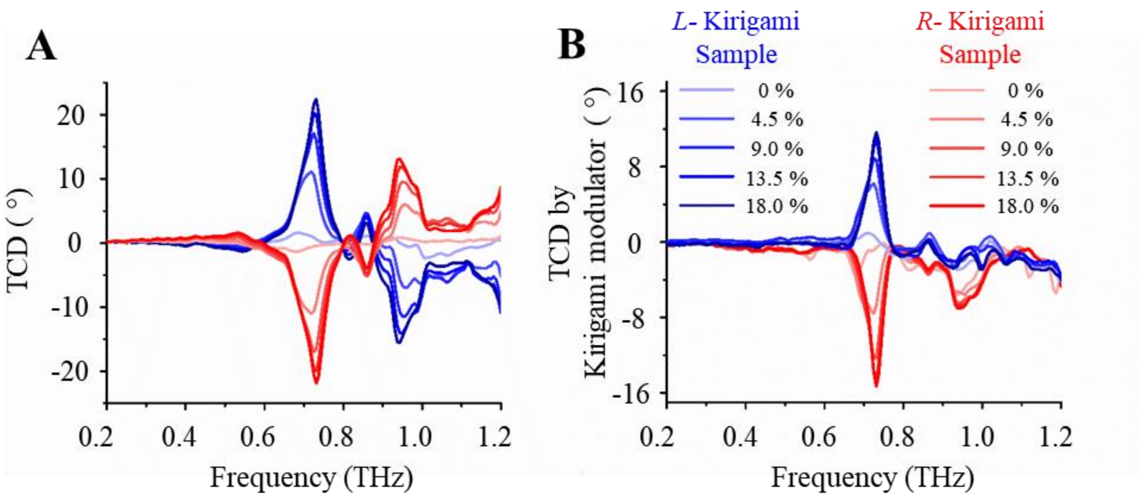


**Fig. S14. TCD spectra of kirigami samples with slant angle of 37.5° using two linear polarizers method and kirigami modulator method.** (A) TCD spectra measured by standard two polarizers method. (B) TCD spectra measured by kirigami chiral modulator. Blue and red curves are for left-handed (*L*) and right-handed (*R*) samples respectively. The strains applied are given in the legends.

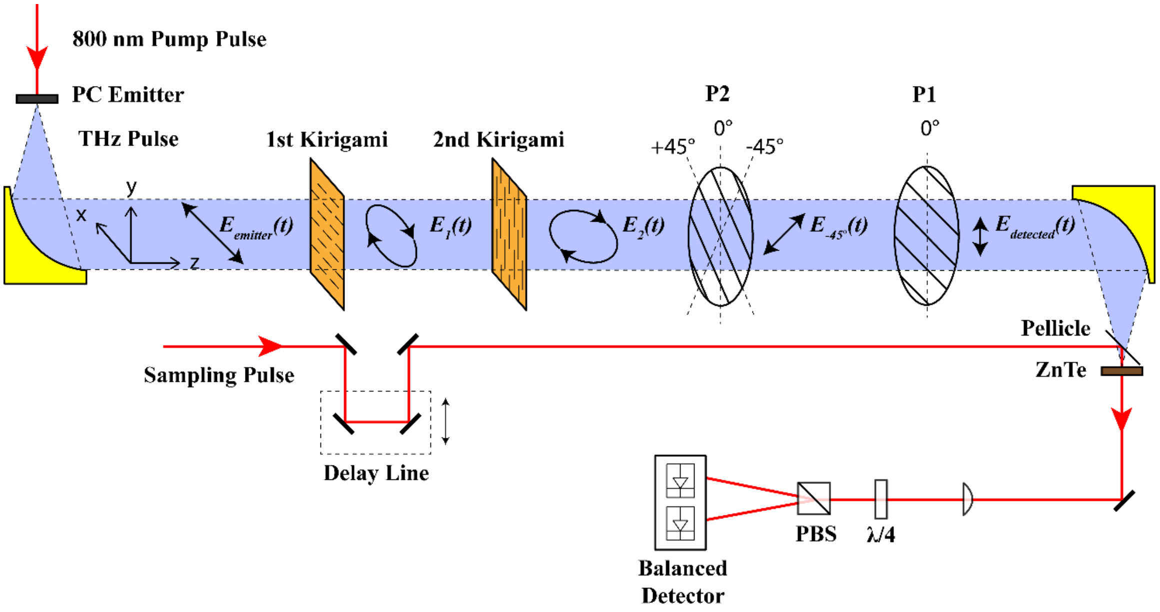


**Fig. S15. Schematic of the experimental setup for stacked kirigami configuration.** A second kirigami sheet is inserted behind the first one to form a double stack configuration and they are controlled by two piezo-controllers, independently. The configuration of the two kirigami sheets shown here is that the first kirigami mounted vertically and the second kirigami mounted horizontally.

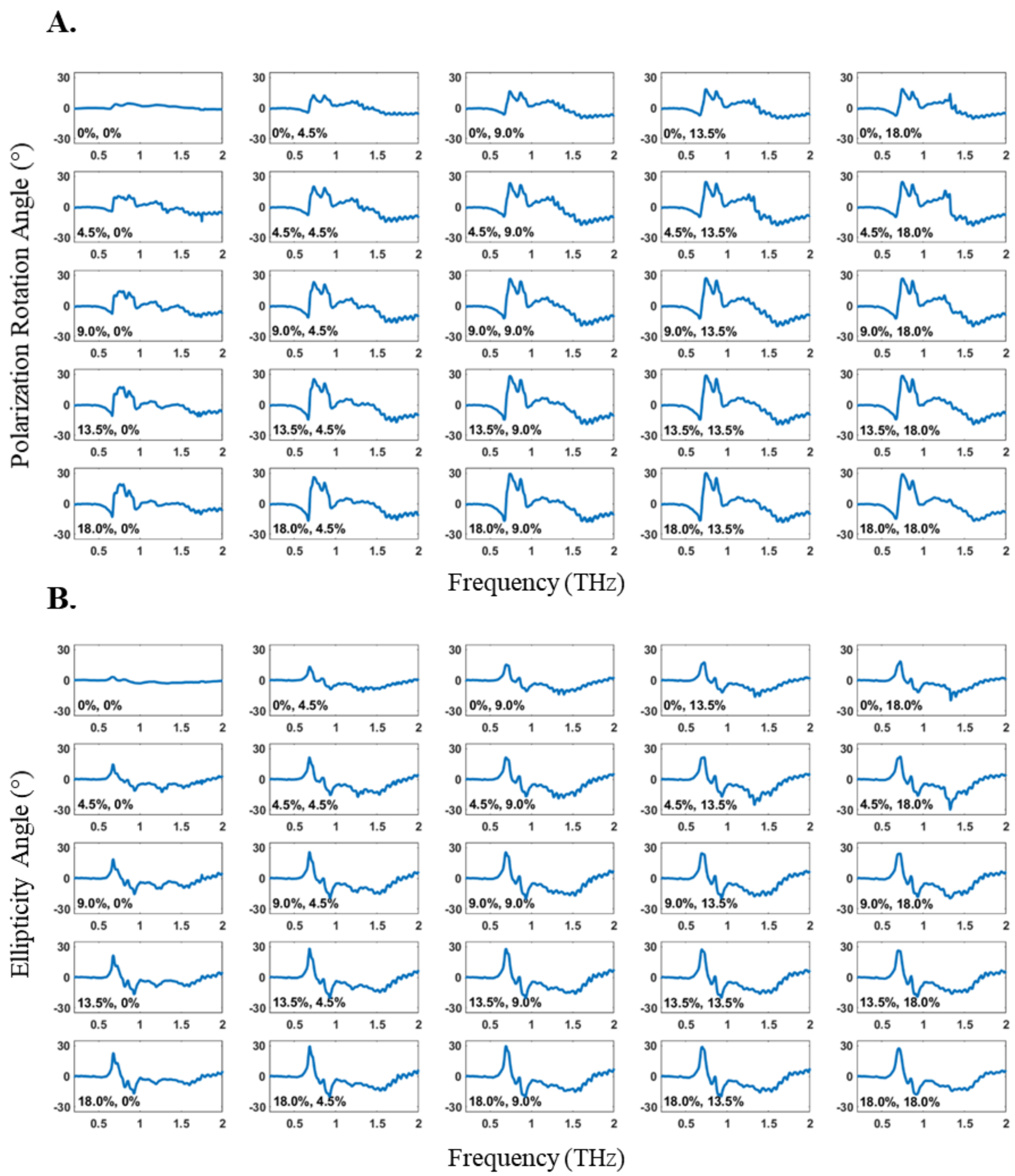


**Fig. S16. Experimental results for vertically-positioned left-handed (VL) kirigami and horizontally-positioned left-handed (HL) kirigami stack.** (A) Polarization rotation angle and (B) ellipticity angle measured from the double stacked kirigami sheets. The first and second number in the legends are the strains applied to the first and second kirigami sheets, respectively. Both kirigami sheets have 37.5° slant angle ($\varphi$).

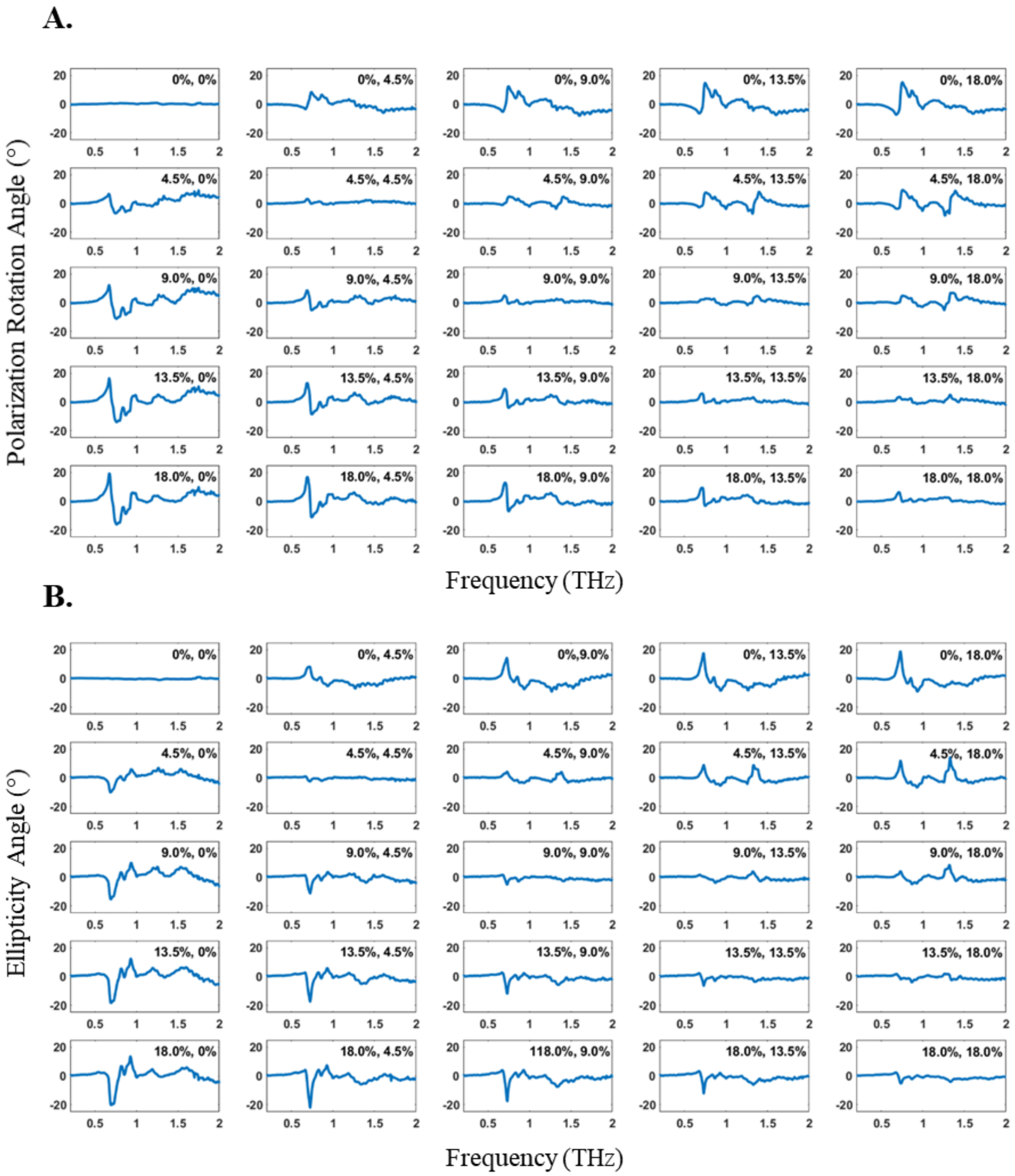


**Fig. S17. Experimental results for vertically-positioned right-handed (VR) kirigami and horizontally-positioned left-handed (HL) kirigami stack.** (A) Polarization rotation angle and (B) ellipticity angle measured from the double stacked kirigami sheets. The first and second number in the legends are the strains applied to the first and second kirigami sheets, respectively. Both kirigami sheets have 37.5° slant angle ($\varphi$).

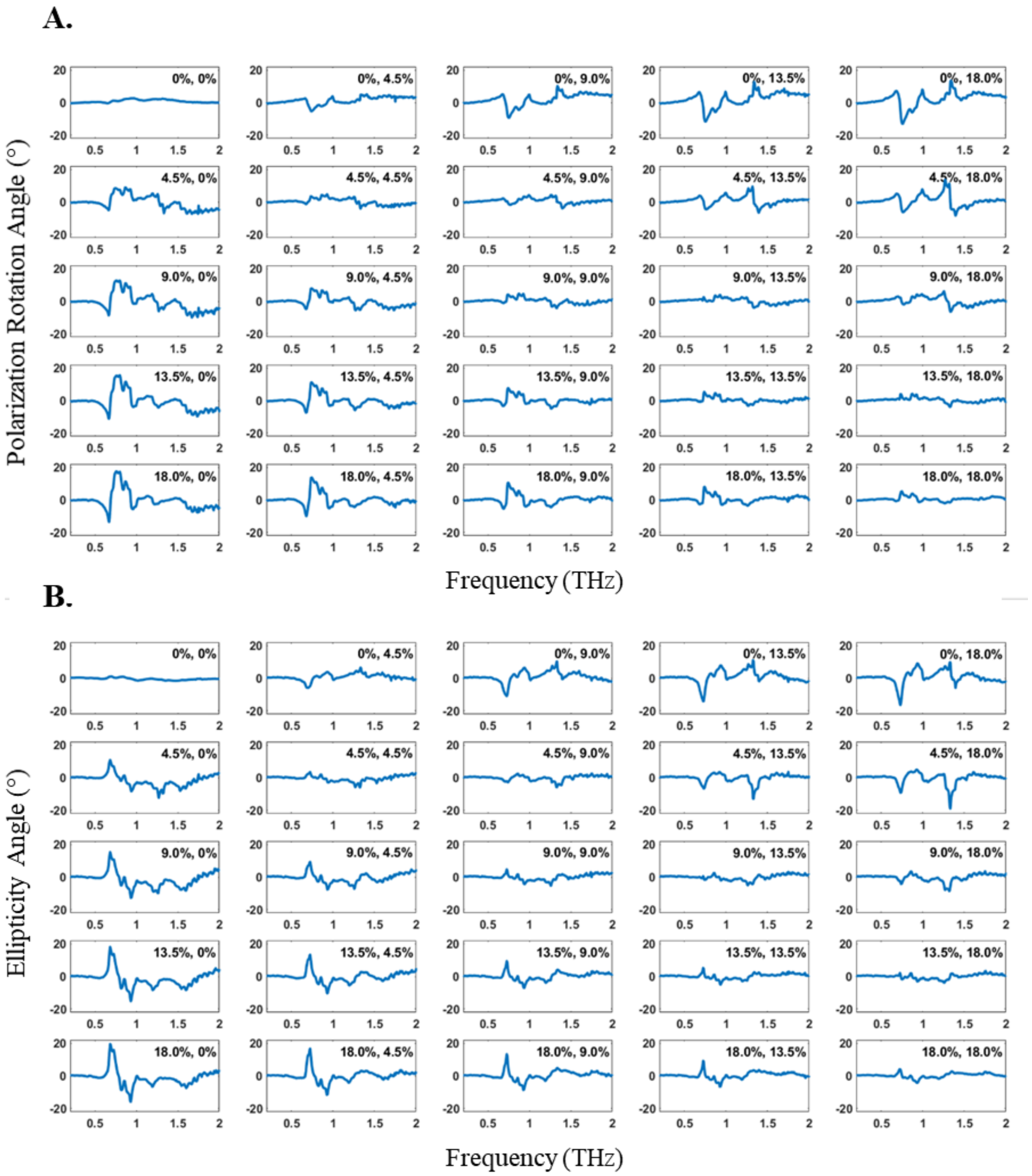


**Fig. S18. Experimental results for vertically-positioned left-handed (VL) kirigami and horizontally-positioned right-handed (HR) kirigami stack.** (A) Polarization rotation angle and (B) ellipticity angle measured from the double stacked kirigami sheets. The first and second number in the legends are the strains applied to the first and second kirigami sheets, respectively. Both kirigami sheets have 37.5° slant angle ($\varphi$).

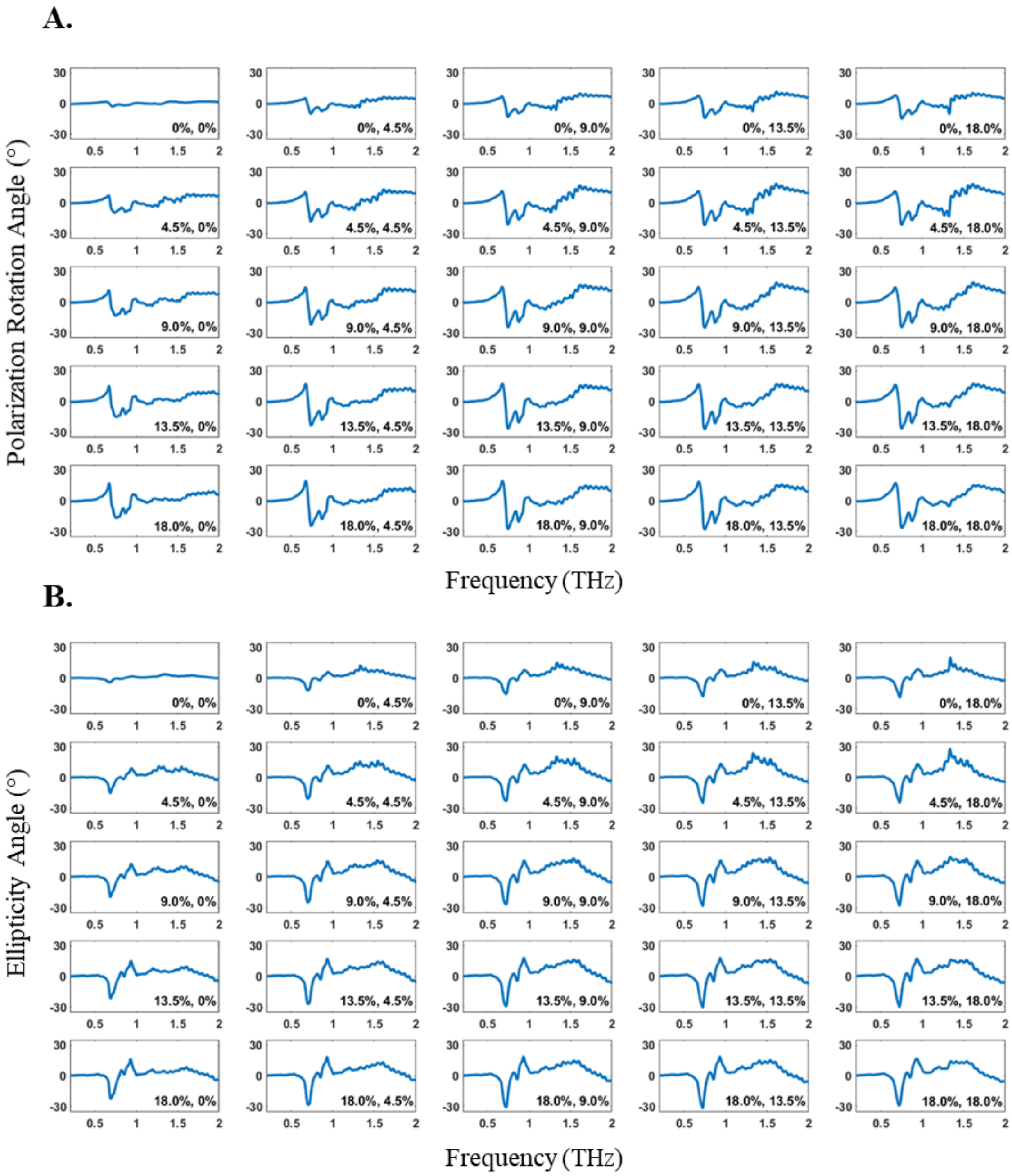


**Fig. S19. Experimental results for vertically-positioned right-handed (VR) kirigami and horizontally-positioned right-handed (HR) kirigami stack.** (A) Polarization rotation angle and (B) ellipticity angle measured from the double stacked kirigami sheets. The first and second number in the legends are the strains applied to the first and second kirigami sheets, respectively. Both kirigami sheets have 37.5° slant angle ($\varphi$).

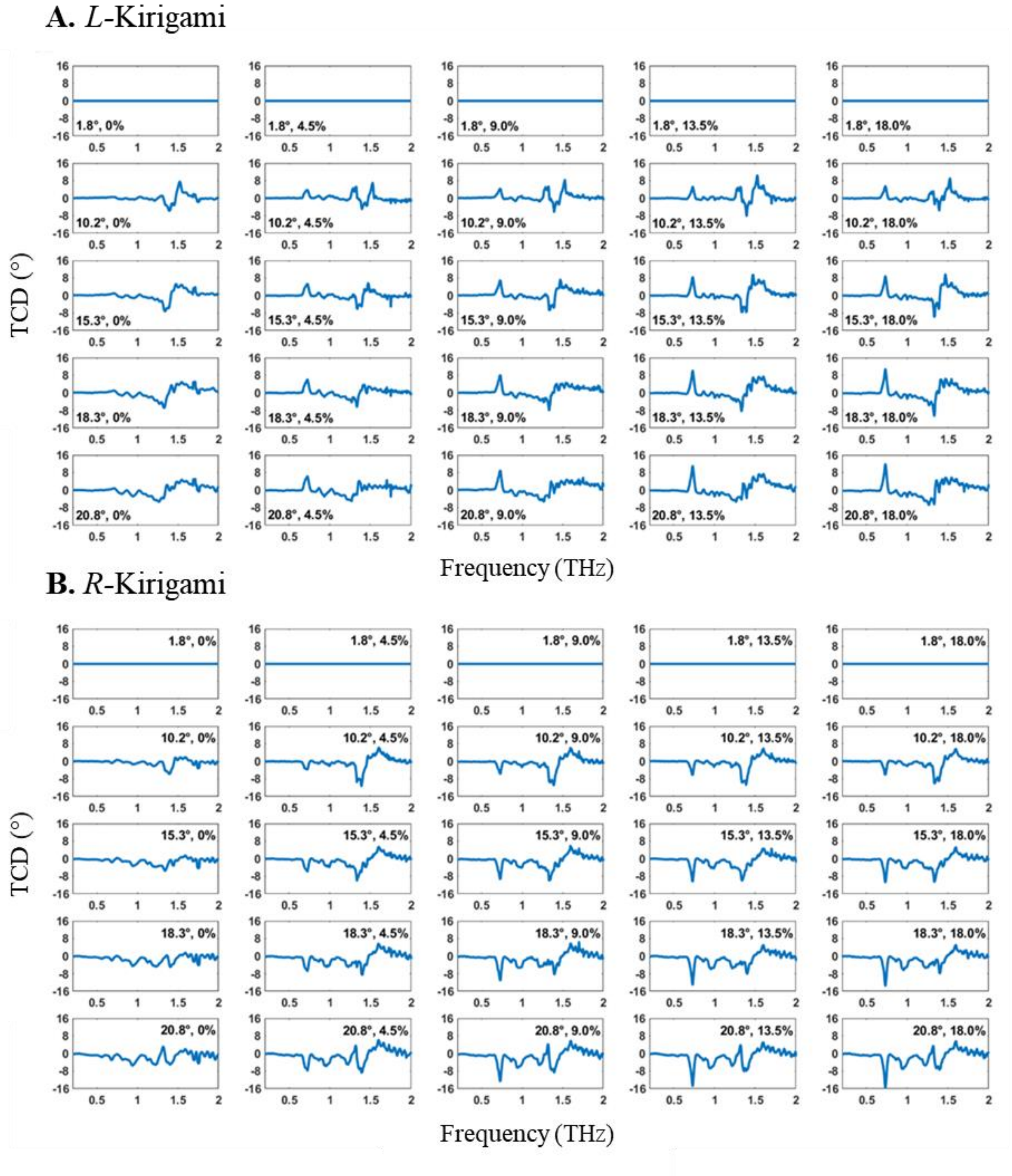


**Fig. S20. Experimental TCD spectra measured by kirigami modulator.** TCD spectra of left-handed kirigami (A) and right-handed kirigami (B) samples with slant angle $\varphi$ of 37.5°. The first number in the legends is the ellipticity of incident THz beam generated by the kirigami modulator (first kirigami) at 0.73 THz. The second number is the strain applied to the second kirigami which controls the chirality of the probed sample.

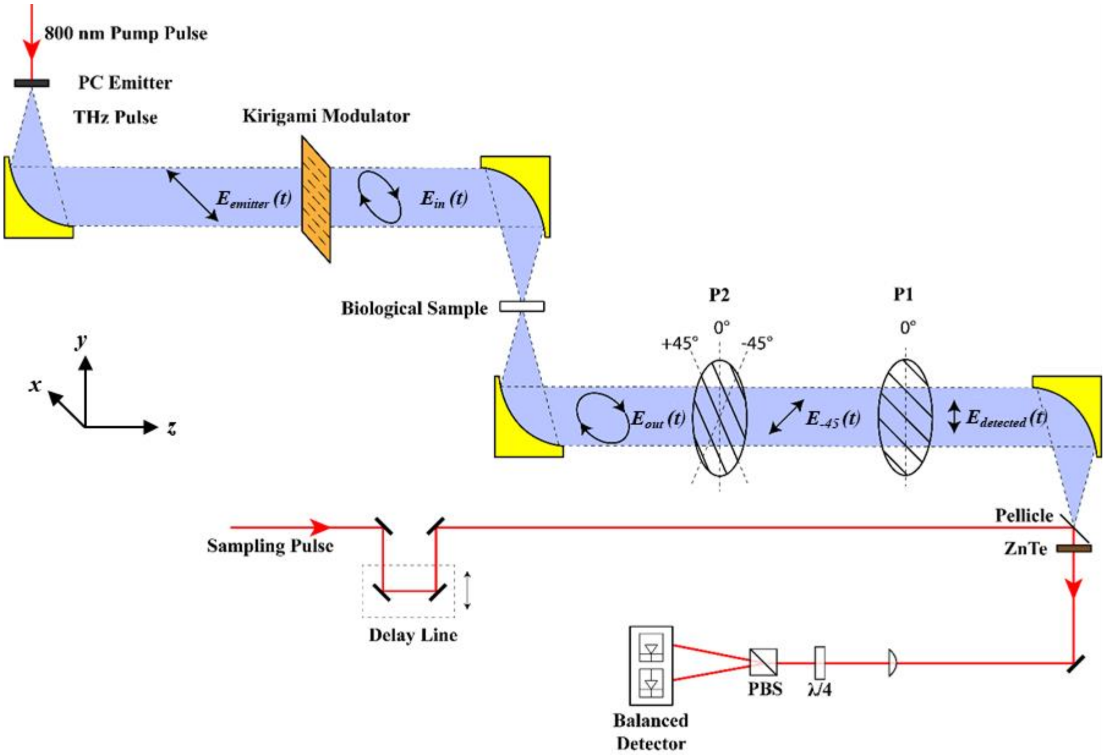


**Fig. S21. Schematic of the experimental TCD setup for biological samples.** The elliptically/circularly polarized THz beam generated by the kirigami modulator is focused by an off-axis parabolic gold mirror to a spot size of approximately 500 μm and acts as the input for the biological samples. The transmitted THz beam through the sample is collected and collimated by another off-axis parabolic gold mirror for detection.

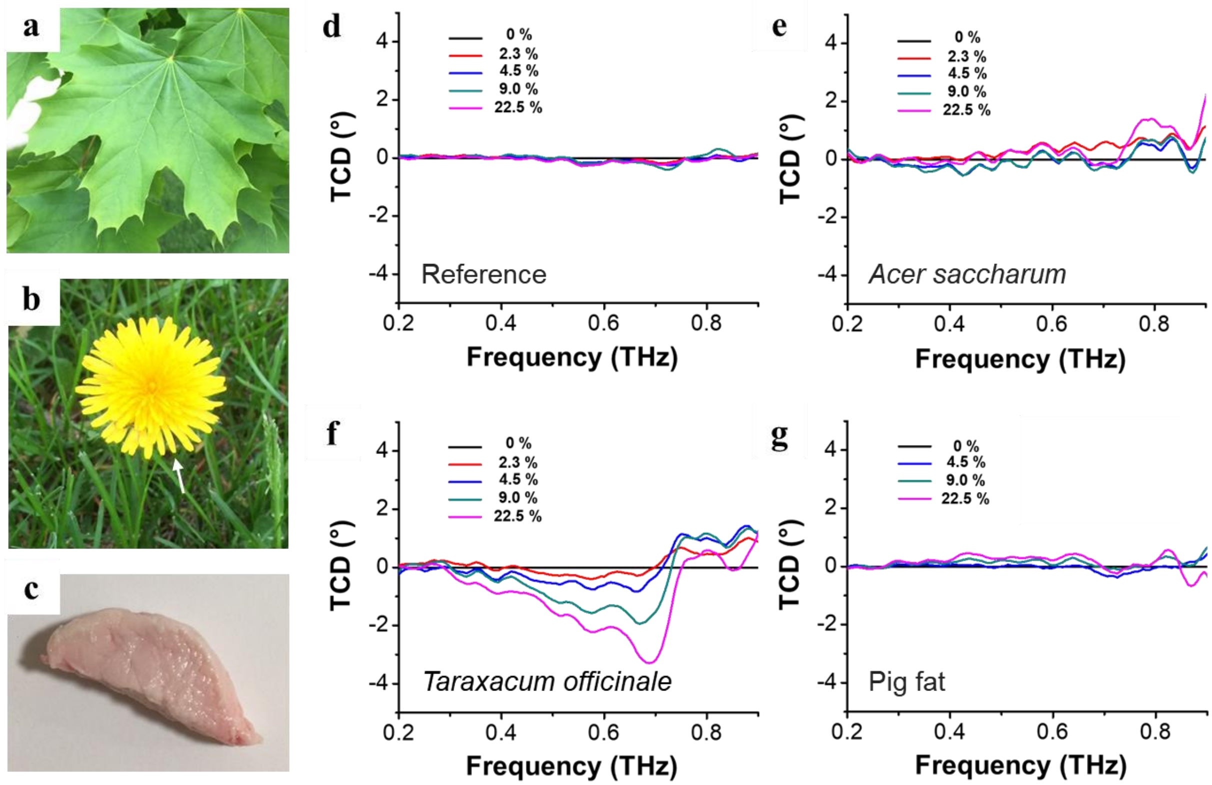


**Fig. S22. Experimental TCD spectra modulated by kirigami of biological samples.** Photographs of a leaf of maple sugar tree (**a**), a petal of dandelion (**b**) and a piece of pig fat (**c**). The arrow in the dandelion image in (**b**) indicates the actual sample for the measurement. (**d**) to (**g**) show TCD spectra of reference, a leaf (*Acer saccharum*), a petal (*Taraxacum officinale*) and a piece of pig fat, respectively. The legend shows the strains applied to the kirigami modulators. The TCD curves for each sample were normalized to its $\varepsilon = 0$ % curve.

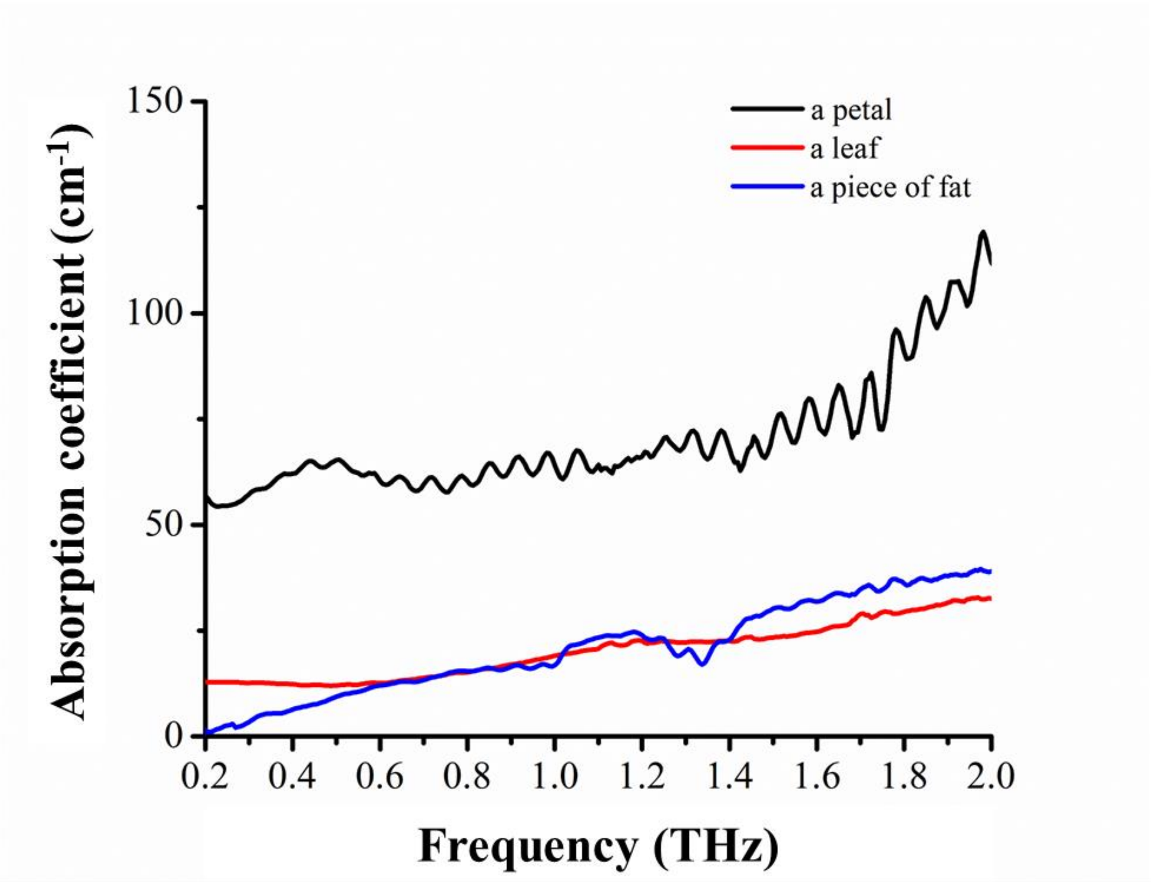


**Fig. S23. Absorption coefficient spectra of three biological samples.** Absorption coefficients of a petal of a dandelion, a leaf of a maple tree and a piece of pig fat. There is a substantial absorption between 0.3 ~ 0.8 THz especially for a petal of a dandelion.

**Movie S1**

Movie of stretching and releasing of kirigami chiroptical modulator ($\varepsilon$ from 0 % to 22.5 %)

**Movie S2**

Movie of 3D topology of a reconstructed kirigami structure with 45° wire slant angle stretched by 13.5% strain.
